# MaAsLin 3: Refining and extending generalized multivariable linear models for meta-omic association discovery

**DOI:** 10.1101/2024.12.13.628459

**Authors:** William A. Nickols, Thomas Kuntz, Jiaxian Shen, Sagun Maharjan, Himel Mallick, Eric A. Franzosa, Kelsey N. Thompson, Jacob T. Nearing, Curtis Huttenhower

**Affiliations:** Department of Biostatistics, Harvard T.H. Chan School of Public Health, Boston, MA, USA; Harvard Chan Microbiome in Public Health Center, Harvard T. H. Chan School of Public Health, Boston, MA, USA; Division of Gastroenterology, Massachusetts General Hospital and Harvard Medical School, Boston, MA, USA; Clinical and Translational Epidemiology Unit, Massachusetts General Hospital and Harvard Medical School, Boston, MA, USA; Division of Biostatistics, Department of Population Health Sciences, Weill Cornell Medicine, Cornell University, New York, NY, USA; Department of Statistics and Data Science, Cornell University, Ithaca, NY; Infectious Disease and Microbiome Program, Broad Institute of MIT and Harvard, Cambridge, MA, USA; Department of Immunology and Infectious Diseases, T.H. Chan School of Public Health, Harvard University, Boston, MA, USA

**Author notes:** Equivalent effort (Kelsey N. Thompson, Jacob T. Nearing, Curtis Huttenhower). Corresponding author: Curtis Huttenhower.

## Abstract

A key question in microbial community analysis is determining which microbial features are associated with community properties such as environmental or health phenotypes. This statistical task is impeded by characteristics of typical microbial community profiling technologies, including sparsity (which can be either technical or biological) and the compositionality imposed by most nucleotide sequencing approaches. Many models have been proposed that focus on how the relative abundance of a feature (e.g. taxon or pathway) relates to one or more covariates. Few of these, however, simultaneously control false discovery rates, achieve reasonable power, incorporate complex modeling terms such as random effects, and also permit assessment of prevalence (presence/absence) associations and absolute abundance associations (when appropriate measurements are available, e.g. qPCR or spike-ins). Here, we introduce MaAsLin 3 (Microbiome Multivariable Associations with Linear Models), a modeling framework that simultaneously identifies both abundance and prevalence relationships in microbiome studies with modern, potentially complex designs. MaAsLin 3 also newly accounts for compositionality with experimental (spike-ins and total microbial load estimation) or computational techniques, and it expands the space of biological hypotheses that can be tested with inference for new covariate types. On a variety of synthetic and real datasets, MaAsLin 3 outperformed current state-of-the-art differential abundance methods in testing and inferring associations from compositional data. When applied to the Inflammatory Bowel Disease Multi-omics Database, MaAsLin 3 corroborated many previously reported microbial associations with the inflammatory bowel diseases, but notably 77% of associations were with feature prevalence rather than abundance. In summary, MaAsLin 3 enables researchers to identify microbiome associations with higher accuracy and more specific association types, especially in complex datasets with multiple covariates and repeated measures.

## Introduction

Quantitative microbial community analyses often aim to identify community features (e.g. microbes, genes, or metabolites) associated with metadata (e.g. human health outcomes, diet, or environmental conditions). Various procedures have been developed to test for these associations, but there is still relatively little consensus on how to best approach differential abundance (DA) testing in microbiome studies.^1^ In large part, this is driven by the unusual properties of microbiome data, including compositionality, sparsity, and heteroskedasticity.^2, 3, 4, 5, 6^ Like most ‘omics data, they also generally present challenges due to their inherent high dimensionality,^7^ requiring efficient algorithms and corrections for multiple hypothesis testing.

In response to these challenges, a variety of methods for identifying differentially abundant features have been developed for microbial community profiles. Though none is perfect, three widely-used examples that produce consistent and accurate benchmarking results are ALDEx2, ANCOM-BC, and MaAsLin 2.^1,8, 9^ To address compositionality in microbiome data, ALDEx2 (ANOVA-Like Differential Expression) updates a Bayesian Dirichlet prior based on the observed read counts, performs a centered log-ratio transformation on samples from the posterior, and runs a significance test on these posterior ratios.^10^ In a different strategy, ANCOM-BC (Analysis of Composition of Microbiomes with Bias Correction)^11^ (and subsequently ANCOM-BC2^12^) uses an expectation maximization procedure to estimate the compositionality-induced bias when comparing abundances between multiple groups; it then tests for differences in the groups after accounting for this bias. Finally, MaAsLin 2 (Microbiome Multivariable Associations with Linear Models) fits linear models on log-transformed relative abundances, as its evaluations found this to be comparably accurate to explicitly accounting for compositionality while preserving flexibility and numerical stability.^13^ Furthermore, several evaluations have found that simple models like MaAsLin’s often outperform more complicated DA methods.^14, 9, 15^

These methods also account for zero inflation with different strategies. In ALDEx2, a pseudocount of 1/2 is used for each taxon in each sample in the Bayesian prior. In ANCOM-BC2, zeros are split into outlier zeros (treated as missing data), structural zeros (taxa that are always zero in one group, marked as differentially abundant), and sampling zeros (replaced with pseudocounts). When pseudo-counts are used, ANCOM-BC2 also tests the model’s sensitivity to the magnitude of the pseudo-count. MaAsLin 2 also uses a pseudo-count for zeros, setting them to half the minimum observed relative abundance per feature. Importantly, apart from ANCOM-BC2’s handling of structural zeros, these methods only test whether feature prevalence (presence or absence) is associated with covariates insofar as the prevalence association contributes to the abundance association. Some more niche methods explicitly separate prevalence and abundance testing,^16, 17^ but they either do not handle multiple covariates or do not give effect sizes in addition to significance tests. However, aside from technical reasons, microbial presence alone is of course phenotypically relevant in many cases, above and beyond abundances: pathogens often cause disease even at extremely low abundance,^18, 19^ some taxa are detected more frequently in e.g. colorectal cancer patients,^20^ and many taxa have only been observed to occur in certain geographic, demographic, or racial/ethnic groups.^21, 22, 23^ Thus, detecting phenotypes associated specifically with the presence or absence of microbial features—hereafter referred to as prevalence differences—would be scientifically advantageous.^24, 9^

To bridge these gaps, MaAsLin 3 (1) explicitly identifies prevalence associations in addition to abundance associations; (2) accounts for compositionality with a median coefficient comparison strategy, or by using data from absolute abundance protocols when available; and (3) extends its previous inference methods to include generalized mixed effects models, group-wise differences, and ordered predictors. It natively handles both count and precomputed relative abundance data, and it can be applied to any high dimensional ‘omics data including taxonomic profiling, functional profiling, and metatranscriptomics. In simulations, MaAsLin 3’s precision improved by up to 0.29 on average over MaAsLin 2 while maintaining the same recall and distinguishing between abundance and prevalence associations, and it performed favorably when compared to other DA methods. When applied to the Human Microbiome Project 2 (HMP2) Inflammatory Bowel Disease Multi-omics Database (IBDMDB),^25^ it identified significant features that largely agreed with previous results, save that 77% of identified associations were prevalence-based. MaAsLin 3, along with its source code, documentation, tutorials, and example data sets are freely available in the MaAsLin 3 R/Bioconductor software package and at https://huttenhower.sph.harvard.edu/maaslin3.

## Results

Here, we present MaAsLin 3, which carries out microbiome feature DA testing while accommodating sparsity, compositionality, high dimensionality, complex experimental designs, experimental protocols that provide absolute abundance information, and phenotypes dependent on microbial feature presence or absence. The last in particular is assessed using a logistic model for presence/absence and a log-linear model for non-zero abundances, and MaAsLin 3 accounts for compositionality in abundance coefficients by comparing coefficients against their median. The MaAsLin 3 implementation takes as input a per-sample feature table and a metadata table, and it returns a table of associations including estimated coefficients and significance values along with summary and per-association plots. Relative to MaAsLin 2, MaAsLin 3 distinguishes between prevalence and abundance associations, accounts for compositionality with experimental or computational techniques, and enables inference for new covariate types. These improvements allow users to identify more specific biological associations, better approximate absolute abundance effects, and account for more complex study designs.

MaAsLin 3’s algorithm consists of (1) normalizing microbial community feature abundance profiles (by default, total-sum scaling to obtain relative abundances), (2) generating a parallel prevalence (present (1) versus absent (0)) profile and retaining a separate subset of data with non-zero abundances, (3) log transforming (base 2) the abundance dataset to stabilize variance, (4) performing a modified logistic regression on the prevalence dataset and linear regression on the abundance dataset, and (5) combining the quantitative effects into an overall effect for each feature-metadatum pair (**Fig. 1a, Methods**). MaAsLin 3 is therefore classified as a hurdle model with a distribution for presence/absence and a distribution for present values, like MAST or scREHurdle for single-cell RNA sequencing.^26, 27^ However, it differs in key ways from other zero-inflated models for microbiome data such as metagenomeSeq^28^ that do not model prevalence associations with covariates.

**Figure 1:**
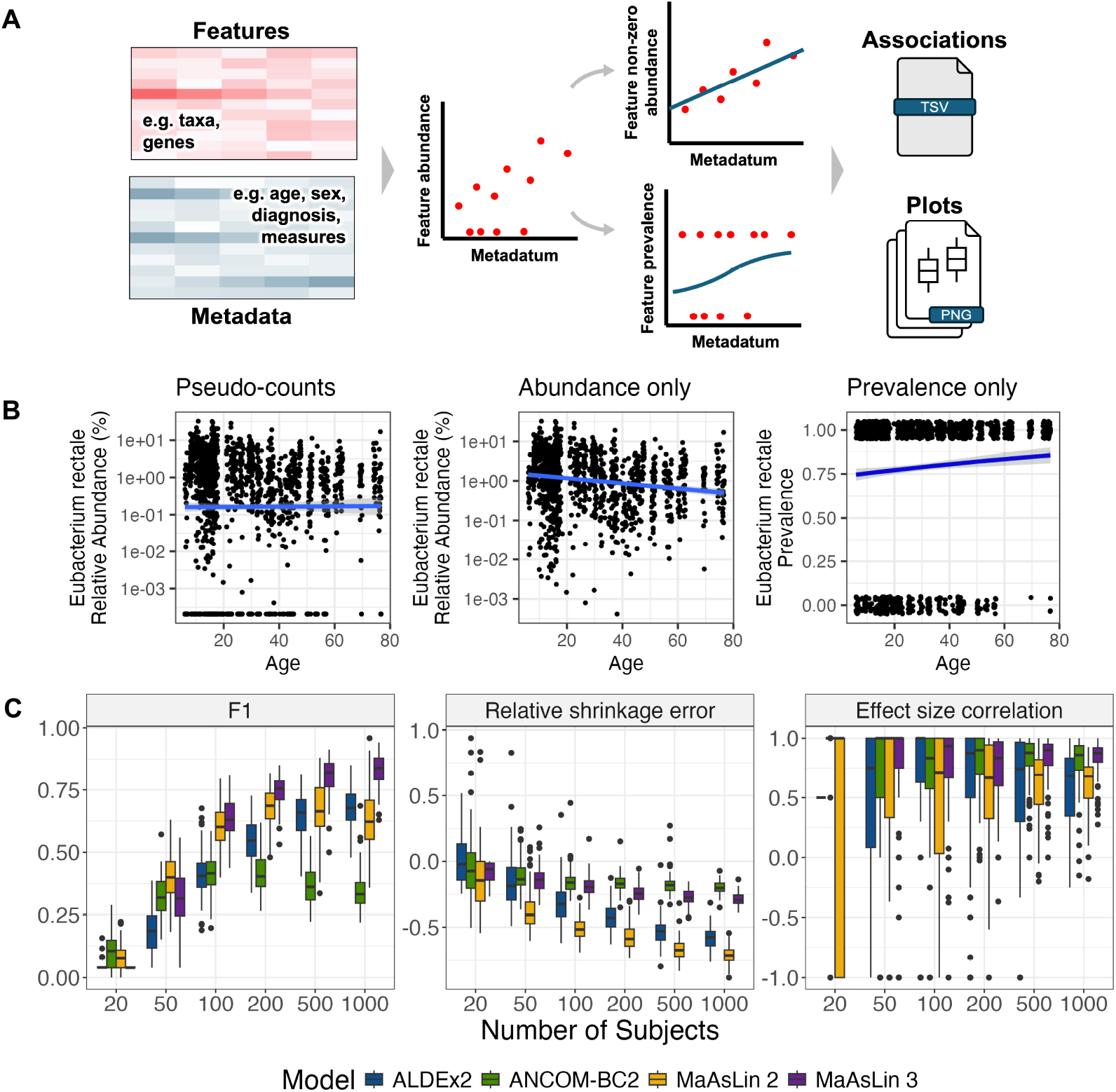
MaAsLin 3 enables both abundance and prevalence modeling with improved accuracy. **A**. MaAsLin 3 model overview. MaAsLin 3 takes as input a table of microbial community feature abundances, as counts or relative abundances, and a corresponding set of metadata (phenotypes, covariates, exposures, etc.). These feature data are normalized, filtered, split into prevalence and log-transformed non-zero abundances, and fit with a modified logistic model or a linear model, respectively. A table of associations is produced indicating the summary statistics corresponding with each feature-metadatum association. **B**. Using all metagenomes from the HMP2 IBDMDB cohort,^25^ *Eubacterium rectale* shows no association with age when zeros are replaced with pseudo-counts, but it shows a negative non-zero abundance association and a positive prevalence association. **C**. MaAsLin 3 out-performs other DA methods in simulations. MaAsLin 3 and other common DA methods were run on 100 synthetic log-normal datasets from SparseDOSSA 2.^5^ For these simulations, 100 features and 5 metadata were simulated with 10% of the feature-metadatum pairs having true associations with coefficients sampled uniformly from 2.5 to 5, half of which were positive and half of which were negative. Half of the true associations were abundance associations; the rest were prevalence associations. The read depth per sample was drawn from a log-normal distribution with a mean of 50,000 (analogous to the number of informative reads per dataset, such as amplicon sequencing). Significant associations (no model fitting errors, q-value less than 0.1, joint q-value for MaAsLin 3) were considered correct if they matched the true associations in the feature and metadatum. A mismatch in association type—abundance versus prevalence—was allowed for all methods since no method besides MaAsLin 3 reports association type. F1 is the harmonic mean of precision and recall; 1 is optimal. The relative shrinkage error is the difference between the absolute fit and true coefficients divided by the true coefficient, averaged over the significant associations; 0 is optimal. The effect size correlation is the Spearman correlation between the fit and true coefficients per metadatum averaged over the metadata; 1 is optimal. Each point represents a simulated dataset.

Separating prevalence modeling—whether a feature is present or absent—from abundance modeling—the quantity of a feature when present—is useful for both technical and biological reasons. Technically, many common microbiome data transformations (e.g. log) require omission or replacement of zeros are with pseudo-counts, but downstream results can be sensitive to this choice of pseudo-count.^29, 12, 16^ Biologically, the presence vs. abundance of a microbe can easily correspond with different phenotypic phenomena, such as pathogen infection versus commensal resource consumption. Additionally, the two effects can sometimes quantitatively cancel: a feature might have a negative abundance association with a metadatum but a positive prevalence association with that metadatum, a combination that causes the association to be undetected when aggregated. As a motivating example, in the IBDMDB cohort, when *Eubacterium rectale* is regressed against age in a simple linear model, there is no association when using pseudo-counts set to half the dataset minimum (*β* = 0.001, *p* = 0.86) (**Fig. 1b**). However, when split into a logistic regression for the presence/absence transformation of the data and a linear regression for the non-zero portion of the data, significant but opposite associations emerge (linear *β* = − 0.015, *p* = 1 *×* 10^*−*9^; logistic *β* = 0.010, *p* = 0.005). An interpretation would be that carriage of *E. rectale* becomes more likely with age, but the amount of *E. rectale* when present tends to decrease. Such examples motivate the separation of abundance and prevalence testing.

In addition to prevalence modeling, another major improvement of MaAsLin 3 is the option to identify associations either on a true (experimental) absolute abundance scale or by including an additional (inferred) component accounting for compositionality. When only testing for relative abundance associations, MaAsLin 3 simply fits the specified models on relative abundance data. Biologically, both relative and absolute abundance phenotypes are plausible, but more sophisticated methods are necessary to provide an option for also inferring potential absolute-scale associations.^2, 3, 4, 30^ To identify such associations, MaAsLin 3 can also test for a difference between each feature’s coefficient and the median of these coefficients (**Methods**). When fewer than half of the community’s features are changing in absolute abundance—a frequent and biologically plausible assumption in microbiome DA testing^31,16, 32^—this modified test can be interpreted as a test against the null hypothesis that an association’s coefficient is zero on an absolute scale^31^ (**Sup. Fig. 1a, Methods**). However, even when this assumption is violated, the ranking of the coefficients on the relative and absolute scales will be identical in the absence of sparsity and will likely be similar even with sparsity.^3^ Therefore, this median comparison can be interpreted as a test for whether a particular feature’s association with a covariate is different from the typical association between features and the covariate on an absolute scale. As shown below, the associated test results align closely with true absolute abundance associations derived from experimental data.

### MaAsLin 3 improves precision and accuracy relative to other differential abundance methods

To evaluate how MaAsLin 3’s prevalence modeling and compositionality correction perform in data with known associations, synthetic microbiome profiles were simulated from a modified zero-inflated log-normal distribution using SparseDOSSA 2.^5^ SparseDOSSA 2 generates an underlying null distribution of features based on real metagenomic data, adds in abundance (log-linear) and prevalence (logistic) associations on a simulated absolute scale, and samples reads to produce resulting counts and normalized relative abundances. Therefore, the simulated data available for DA testing are on the relative scale, but the underlying associations are known on an absolute scale. The resulting data were analyzed with MaAsLin 3, ALDEx2, ANCOM-BC2, and MaAsLin 2 (all using predominantly default parameters; see **Methods**).

When comparing modeled vs. true associations, MaAsLin 3 obtained a median F1 score as high or higher than any other method on datasets containing more than 50 samples (**Fig. 1C**). MaAsLin 3 and ALDEx2 maintained the highest precision at all sample sizes (average precision *≥* 0.82 and *≥* 0.75), though the precision of all methods decreased with larger sample sizes due to increasing power to discover spurious compositionality-induced associations (**Sup. Fig. 2**). When just considering true prevalence associations in well-powered 1,000 sample settings, even previous methods that did not explicitly model prevalence achieved substantial recall (0.78 for ALDEx2, 0.54 for ANCOM-BC2, and 0.94 for MaAsLin 2). This reinforces that other methods identify prevalence associations as abundance associations since they can be considered a subtype of abundance association. At small sample sizes, the recall of MaAsLin 3 was lower (average 0.19 with 50 samples) than of ANCOM-BC2 (0.23) or MaAsLin 2 (0.27) due to logistic models typically requiring more samples to reach significance and MaAsLin 3’s linear model component only considering non-zero values (reducing effective sample size). However, because significant associations are often targets for time-intensive follow-up analysis, precision is typically more important than recall, and low-precision methodologies can raise concerns about reproducibility.^33^ In these low sample settings, MaAsLin 3 had higher precision (average 0.99 at 50 samples) than the methods with higher recall (0.89 for MaAsLin 2 and 0.54 for ANCOM-BC2).

When comparing the modeled coefficients from each method to the true abundance coefficients, all methods produced coefficients biased towards zero, but MaAsLin 3 and ANCOM-BC2 yielded the least shrinkage (**Fig. 1C**). When methods impute zeros with simple pseudocounts, the fit coefficients become a combination of the true effect and a null effect (zero slope) due to high random sparsity in the data. Despite the shrinkage, with at least 50 samples, all methods produced coefficients that showed moderate to high Spearman correlations with the true abundance coefficients (averages 0.46-0.83), though MaAsLin 3 and ANCOM-BC2 performed slightly better than the rest at high sample sizes (average 0.83 and 0.80 versus 0.62 and 0.59 for MaAsLin 2 and ALDEx2 at 1000 samples) (**Fig. 1C**). Thus, though the coefficients are biased towards zero in models that rely heavily on pseudocounts, the ordering of the effect sizes is generally accurate.

When varying the effect size of the true coefficients, MaAsLin 3 and ALDEx2 retained the highest precision across all effect sizes (average precision *≥*0.86 and *≥*0.88), with MaAsLin 2 losing precision at high effect sizes and ANCOM-BC2 having low precision at all effect sizes (average precision *≤* 0.52; **Sup. Fig. 3**). ANCOM-BC2 had slightly higher recall at very low effect sizes (average 0.10 with effect sizes 0.5-1), but MaAsLin 2 and MaAsLin 3 had the highest recall with effect sizes above 2.5. Coefficients fit by the models were shrunk more when the true coefficients were larger, and, for each method, the correlation between the fit and true coefficients was similar for any true effects larger than 1.

When multiple observations per subject were available, the precisions of ALDEx2, MaAsLin 2, and MaAsLin 3 all decreased with more samples per subject, as the models had higher power to discover spurious compositional effects. The precision decrease was slightly larger for MaAsLin 3 (0.97 with no repeats versus 0.75 with 10 samples per subject) than for ALDEx2 (0.97 versus 0.80), but MaAsLin 3 demonstrated much higher recall (average 0.80 at 10 samples per subject versus 0.55 for ALDEx2). Most of MaAsLin 3’s precision loss was attributable to small but significant prevalence effects induced by compositionality: on average, 70% of false discoveries at 10 samples per subject involved fit prevalence coefficients with absolute values less than 1. Therefore, a post-hoc threshold of 1 was applied to the coefficients from all models (i.e., only associations with absolute effects greater than 1 were evaluated; **Sup. Fig. 4B**). This returned the precision of MaAsLin 3 to a high level (average *≥* 0.92 for any repeat number) similar to ALDEx2 and ANCOM-BC2 without affecting the recall. In contrast to all of these results, the precision of ANCOM-BC2 increased substantially with multiple observations (average 0.52 with no repeats versus 0.99 with 2 samples per subject), though its recall remained low (average *≤* 0.33 for any number of repeats; **Sup. Fig. 4B**). All coefficients correlated well with their true values, though MaAsLin 3 and ANCOM-BC2 correlated better (average 0.82 and 0.84 at 10 repeats versus

0.54 and 0.62 for ALDEx2 and MaAsLin 2). Taken together, this suggests that MaAsLin 3 accurately identifies associations in longitudinal data, and precision can be further improved by applying a small threshold to the coefficients (e.g. set to 20% of the largest reliable coefficients).

### MaAsLin 3 is robust to violations of modeling assumptions

To evaluate MaAsLin 3’s performance on data not conforming to its model assumptions (i.e. log-normal abundances with logistic prevalence associations), we used the evaluation procedure of ANCOM-BC^11^ to simulate datasets with added abundance effects and structural zeros in which all samples from one group lack the feature (**Methods**). By design, the ANCOM-BC simulations are somewhat pathological with respect to both sparsity and compositionality. With these data, only MaAsLin 3 and ANCOM-BC2 maintained high precision at low sample sizes (average precisions 0.96 and 0.96 at 50 samples), though both had substantial precision losses above 100 samples, with ANCOM-BC2 losing slightly more precision (at 1,000 samples, average precision 0.64 for MaAsLin 3 and 0.59 for ANCOM-BC2; **Sup. Fig. 5**). At any sample size, the average precision of ALDEx2 and MaAsLin 2 never exceeded 0.49. All methods except ALDEx2 produced nearly perfect recall with at least 50 samples, and the fitted effect sizes correlated well with true effect sizes for all methods at all sample sizes (all average correlations *≥* 0.92). MaAsLin 3 and ANCOM-BC2 produced slightly shrunk coefficients (average −12% and −13% at 1000 samples), while MaAsLin 2 produced increasingly inflated coefficients (average 7.9% at 1000 samples), and ALDEx2 produced approximately unbiased coefficients with at least 100 samples. Thus, MaAsLin 3 maintained nearly as good or better accuracy than other methods even when its assumptions were violated.

As an additional check of robustness, we recreated a previously published randomization procedure^1^ by randomizing (100 times) the metadata from 38 datasets with binary meta-data and running MaAsLin 3 on the corresponding feature abundance tables. In every replication, fewer than 1% of joint q-values were significant, and most replications (90%) had no significant joint q-values at all (**Sup. Fig. 6**). When splitting by association type, fewer than 1% of the abundance q-values were significant for any replication, and only 9 of the 3,800 replications produced any false positive prevalence associations. These results are similar to the corresponding previous analysis on ALDEx2, ANCOM-II, and MaAsLin 2.^1^ By contrast, when run on the original (non-randomized) datasets, MaAsLin 3 identified many associations in most datasets: on average, 2.3% of abundance associations were significant, 3.3% of prevalence associations were significant, and 6.5% of overall associations (combining prevalence and abundance q-values per association, **Methods**) were significant. This also suggests that prevalence associations might often be more common than abundance associations in real data, for reasons biological (e.g. niche specialization or individual personalization), statistical (e.g. only associations large enough to involve feature loss can be detected), or technical (e.g. sequencing depth restricts detecting abundance changes).

### MaAsLin 3’s model components improve accuracy beyond simpler regression frameworks

We next sought to understand which features of MaAsLin 3’s modeling strategy contributed to improved accuracy by evaluating the abundance and prevalence associations separately. In both a 100-sample typical scenario and a 1,000-sample high-power scenario designed to expose flaws in the model, MaAsLin 3’s default median abundance comparison approach improved precision for any fixed recall level (**Fig. 2A**). To test the inference of associations from only relative abundances against a model incorporating absolute abundance information, we modified SparseDOSSA 2 to mimic a spike-in experiment in which a known quantity of a reference feature is added to each sample and used for normalization (**Methods**). Running MaAsLin 3 on this spike-in data improved precision even further for any fixed recall level, both at 100 and at 1,000 samples. When applying a q-value threshold of 0.1, the default MaAsLin 3 abundance model achieved higher precision than MaAsLin 3 with spike-in normalization (average precision 0.85 versus 0.73 at 1,000 samples) despite the former not requiring experimental modification, at least in settings with relatively few true associations (**Sup. Fig. 7A**). Note that this precision is lower than in the comparison above among DA methods, since here a MaAsLin 3 result was only counted as correct if it also correctly specified whether the association was with prevalence or abundance. As expected, this precision difference was balanced by slightly higher recall for the spike-in normalization strategy (recall 0.75 without versus 0.82 with).

**Figure 2:**
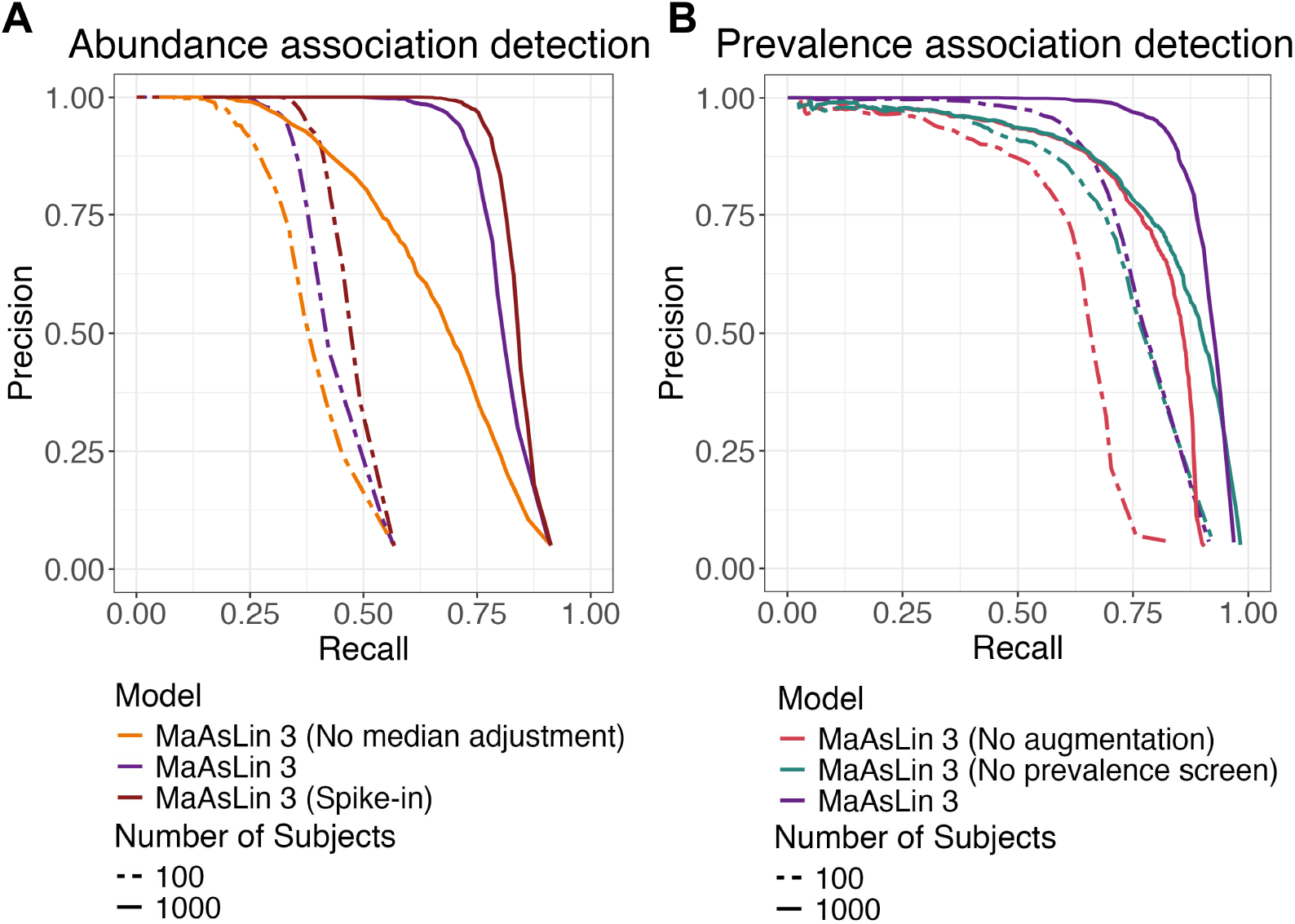
MaAsLin 3’s default model components improve accuracy beyond simpler regression models. **A**. Precision versus recall across a range of q-value thresholds for various MaAsLin 3 abundance modeling options: without a median adjustment for compositionality when only using relative abundance data; MaAsLin 3’s default settings (with the median adjustment when using relative abundance data); and without the median adjustment using data from a (simulated) experimental spike-in procedure. The versions were run on the same 100 synthetic log-normal datasets from Sparse-DOSSA 2 as in **Fig. 1C**. Unlike Fig. 1, significant associations (no model fitting errors, individual q-value less than 0.1) were only considered correct if they matched the true associations in the feature, metadatum, and type of association (prevalence/abundance). **B**. Precision versus recall for MaAsLin 3 prevalence modeling options: without data augmentation to account for separability but with prevalence coefficient screening; without prevalence coefficient screening (allowing any significant prevalence associations) but with data augmentation; and MaAsLin 3’s default setting (with both augmentation and prevalence coefficient screening). The same datasets as in A were used. Curves farther to the right are better.

When evaluating MaAsLin 3’s prevalence modeling, two model properties substantially improved precision over simple logistic regression. First, MaAsLin 3, by default, performs a data augmentation step for prevalence modeling to help address cases of complete separability. For each sample, two additional data points are added: one with the same covariates and the feature set to present, and one with the same covariates and the feature set to absent, a procedure equivalent to the Bayesian Diaconis-Ylvisaker prior that extends directly to mixed effects models (**Methods**).^34, 35^ The user-specified logistic regression is then performed on the augmented dataset but with these augmented data points being given a much lower weight than the original data points (**Methods**). Holding recall constant, this augmentation procedure improves precision over the equivalent simple logistic regression and prevents spurious effects that would result from just a few samples with influential covariates having present or absent taxa (**Fig. 2B**). Second, MaAsLin 3, by default, compares the prevalence coefficient for an association to the corresponding abundance coefficient and flags whether the prevalence effect is likely induced by abundance changes (**Methods**). Because sequencing depth is not infinite, when a feature’s abundance is reduced biologically with a covariate, its likelihood of not being detected increases, especially for rare features, a phenomenon that can manifest as a prevalence effect. Discarding prevalence associations flagged as likely abundance induced also improved precision while holding recall fixed (**Fig. 2B**). Thus, MaAsLin 3’s new default features improve accuracy when detecting both abundance and prevalence associations.

### Remaining inaccuracies in MaAsLin 3, in contrast to other models, are attributable to finite read depth

Despite the improvements of MaAsLin 3 over previous methods and simpler regressions, precision and recall remained imperfect at high sample sizes, even under otherwise ideal simulations (**Sup. Fig. 7A**). To investigate the remaining error, we increased the read depth from 50,000 to 50,000,000 reads (in such simulations, each read is equally informative, akin to a marker gene sequencing read). While this ultra-deep read depth is impractical in e.g. most 16S experiments, it serves to demonstrate that the remaining inaccuracies in MaAsLin 3, at least in well-characterized communities, are due to the limitations of finite read depth. In this ultra-deep sequencing scenario, the precision of the MaAsLin 3 prevalence modeling and the abundance modeling with spike-in normalization increased to their nominal 0.9 level (average precision 0.94 and 0.90; **Sup. Fig. 7B**). The recall of the prevalence and abundance models (both with and without spike-in normalization) increased to nearly 1 (all models above 0.87 on average); the relative shrinkage error was almost eliminated (all shrinkages at worst −8.2% on average); and the Spearman correlations increased to nearly 1 (all at least 0.87 on average). These improvements reflect the dual benefits of (1) reducing variability in the observed feature abundance due to variability in which reads happen to be sequenced and (2) reducing the probability that a feature truly present will not be detected due to none of its reads being sequenced. However, these improvements were not as consistent or substantial for other DA methods (**Sup. Fig. 7C**).

### MaAsLin 3 accurately approximates absolute associations using relative abundances, subject to inherent limitations

Until now, all simulation results assumed that only a small proportion (10%) of the feature-metadatum associations were non-null, an assumption common in DA methods.^31,16^ However, this assumption can be violated in practice if most features in the community are changing in abundance. To demonstrate that no method reliant on only relative abundances is exempt from this assumption, SparseDOSSA 2 was used to create datasets in which increasing proportions of the features were positively associated with the metadata on the absolute scale. According to compositional data theory,^3^ the relative abundance coefficients may be biased from the absolute abundance coefficients by a per-metadatum additive shift, but they should correlate perfectly (or, given sparsity, at least correlate well). As expected, the inferred coefficients from all relative abundance models were negatively biased. This bias became more extreme as more features were positively associated with the metadata (**Fig 3A**). ANCOM-BC2 and MaAsLin 3 were most robust to the assumption violation, but even they were negatively biased at any level and very negatively biased (average bias −2.7 and −3.1) when most features were associated with the covariates. As expected, only MaAsLin 3 with spike-in (i.e. experimental) normalization was fully robust to the assumption violation. By contrast, the correlation between the true abundance effects and the approximated effects was mostly unaffected by the proportion of true associations for all methods.

**Figure 3:**
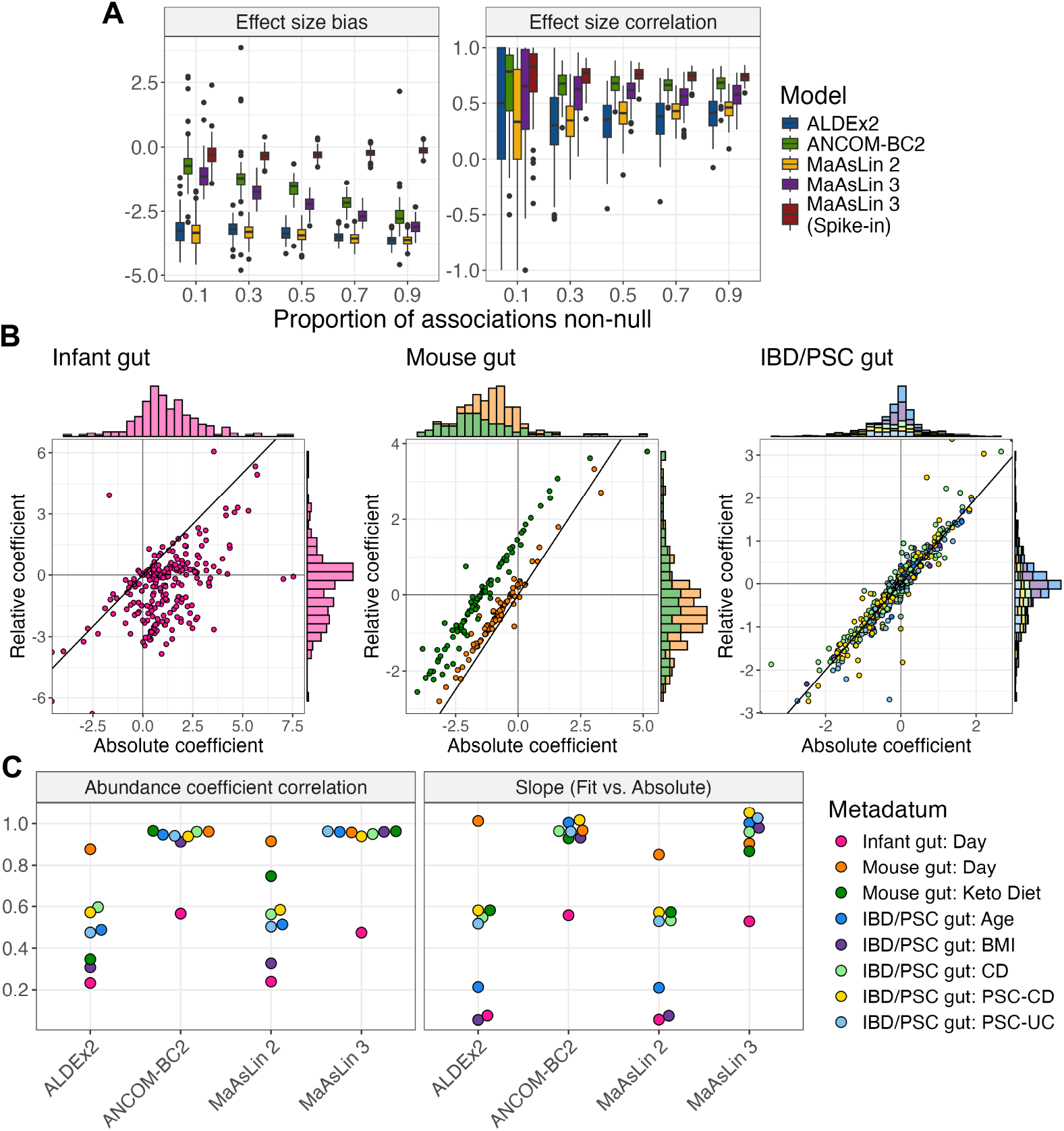
Properties of absolute abundance data that are identifiable on the relative scale are well-identified by MaAsLin 3. A. All methods show increasing bias but little change in coefficient correlation when relying on relative abundance data in which more features have true positive associations. MaAsLin 3 and other common DA methods were run on 100 synthetic log-normal datasets from SparseDOSSA 2 generated as in **Fig. 1C** but with 90% of the associations positive and the sample number fixed at 100. The effect size bias is the mean of the fit coefficients minus their true coefficients for true associations; 0 is optimal. The effect size correlation is the Spearman correlation between the fit and true coefficients per metadatum averaged over the metadata; 1 is optimal. Each point represents a simulated dataset. **B**. Relative and experimentally estimated absolute abundance coefficients agree to varying degrees on three real datasets with experimentally determined (spike-in, digital PCR, or flow cytometry) absolute abundances. MaAsLin 3 was run on both the experimental absolute abundances and on the corresponding relative abundances, and the corresponding coefficients are plotted against each other with one point per feature-metadatum pair. **C**. MaAsLin 3 and ANCOM-BC2 relative abundance regressions best agree with the experimental absolute abundance regressions. ALDEx2, ANCOM-BC2, MaAsLin 2, and MaAsLin 3 were run on the relative abundances and compared to the experimentally determined absolute abundance associations from MaAsLin 3. For each method, for each metadatum in each dataset, the Spearman correlation between the fit relative abundance coefficients and the experimental absolute abundance coefficients was computed over all features. Similarly, for each method, for each metadatum in each dataset, the per-feature relative abundance coefficients were regressed on the per-feature experimentally estimated absolute abundance coefficient. A correlation of 1 and a slope of 1 are optimal.

To evaluate how well these conclusions hold in real data, we re-processed three datasets from diverse communities with absolute abundances experimentally quantified via spikeins, digital PCR, or flow cytometry: an infant gut dataset,^36^ a mouse diet dataset,^37^ and an inflammatory bowel disease and primary sclerosing cholangitis (IBD/PSC) dataset.^38^ Since relative and absolute abundance coefficients only correlate perfectly in the absence of sparsity, these datasets were also chosen for their range of sparsity: the infant dataset contained 97% zeros in its data matrix compared to 66% in the mouse diet dataset and 80% in the IBD/PSC dataset. For each dataset, MaAsLin 3 was run on both the relative and experimentally estimated absolute abundance data, and the corresponding coefficients were compared. As expected, in the sparse infant gut dataset, the relative abundance and experimentally estimated absolute abundance coefficients correlated the least (Spearman correlation 0.47), and the majority of the experimentally determined absolute abundance coefficients were positive (i.e. the taxon increased with time) while a majority of the relative abundance coefficients were negative (**Fig. 3B**). In the mouse diet dataset, the relative and experimentally estimated absolute coefficients were well correlated for both the time variable and the diet variable. However, while the relative and absolute abundance coefficients for time were approximately equal, the diet coefficients showed a clear offset, consistent with relative abundance coefficients being correlated with but shifted from the abundance coefficients. In the IBD/PSC dataset, all relative abundance coefficients were well correlated with their experimentally estimated absolute abundance counterparts, and no systematic shift was observed for any coefficient.

Next, the relative abundance coefficients produced by each method were compared to the experimentally estimated absolute abundance coefficients from MaAsLin 3. Except for in cases of heavy sparsity, a regression of the relative abundances on the absolute abundances should give a slope of 1 and a high correlation. Indeed, for all datasets except the infant gut cohort, MaAsLin 3 and ANCOM-BC2 achieved these results (**Fig. 3C**). By contrast, both ALDEx2 and MaAsLin 2—heavily reliant on pseudo-counts that can bias the abundance relationship—produced coefficients with much weaker correlations (average correlation for non-infant datasets 0.52 and 0.59 versus 0.95 and 0.96 for ANCOM-BC2 and MaAsLin 3) and more attenuated slopes (average slope for non-infant datasets 0.50 and 0.48 versus 0.97 and 0.97). Typically, the infant gut cohort produced both the weakest associations and the most attenuated slopes, suggesting that the assumption of independence between the absolute abundance of the total community and the particular taxa present was severely violated. Indeed, the infant gut is a low-biomass environment, and *Klebsiella* and *Escherichia* blooms were reported in this cohort, reinforcing that the total community absolute abundance depended on the taxa present.^39, 36^ This highlights the importance of using absolute abundance experimental protocols when the community’s total abundance is expected to shift dramatically since both assumptions—half of the features remaining unchanged and independence between the total absolute abundance and the particular taxa present—are violated.

### MaAsLin 3’s linear model extensions enable new experimental designs

In addition to its substantial model extensions to differentiate prevalence and abundance associations, MaAsLin 3 enables five other new types of inference (**Table 1**). First, expanding on the random intercept component of MaAsLin 2, MaAsLin 3 allows the specification of general mixed effects models, including those with interaction terms among metadata covariates. Second, MaAsLin 3 allows testing for omnibus differences among three or more levels of a covariate, expanding categorical testing protocols beyond binary comparisons (**Methods**). When evaluated on SparseDOSSA 2 synthetic data (**Methods**), the omnibus test shows moderate precision (average precision *≥* 0.69) across sample sizes ranging from 20 to 500, with higher precision in the most common 50-100 single study sample range (**Sup. Fig. 8A**). Third, MaAsLin 3 enables testing for level-versus-level differences in ordered predictors, such as dietary frequency data or disease progression (**Methods**). Again when evaluated using SparseDOSSA 2, with sample sizes of 50 or larger, this ordered predictor option maintained high precision (average *≥* 0.88; **Sup. Fig. 8B**). The difference in the precision can be attributed to the fact that the omnibus test does not explicitly account for compositionality, while the ordered predictor option does using the median comparison strategy as above. Fourth, MaAsLin 3 can perform contrast tests among the fit coefficients, which can be useful when testing all pairwise differences among categorical variables (**Methods**). Fifth, MaAsLin 3 natively allows the specification of a feature-specific covariate, such as gene DNA abundance when regressing gene RNA abundance, an important control in metatranscriptomics experiments.^40^

**Table 1:** Linear model extensions enable new experimental designs.

| Feature | Explanation | Example use case |
| --- | --- | --- |
| General mixed model specification | Any valid lme4<sup>41</sup> formula can be specified for the relationship between abundance or prevalence and covariates. | In a study with two treatments and two populations, an interaction term can be specified to evaluate the difference in the treatment’s effect on species abundances between the populations. |
| Omnibus testing | For a covariate with multiple levels (categories), this is an ANOVA-style test of whether the different levels have significantly different coefficients. | In a study with participants from multiple countries, this could test whether species abundances differ among countries without testing for any individual pairwise differences. |
| Level-versus-level differences | For a covariate with ordered levels, this is a test for whether the coefficients for the differences between consecutive levels are significantly non-zero. | In a study of participants with different stages of colorectal cancer, this could test whether species abundances differ between consecutive stages. |
| Contrast testing | This is a test of whether a linear combination of coefficients is significantly different from a constant. When the linear combination is one coefficient minus another and the constant is zero, this tests for a difference in the coefficients. | In a study with samples from different soil types, this could test whether species abundances differ between each pair of soil types. |
| Feature-specific covariates | For a covariate that should be different for each feature, this includes the relevant covariate in each feature’s model. | In any metatranscriptomic study with paired metagenomic data, this controls for a gene’s DNA abundance when regressing that gene’s RNA abundance to identify transcriptional rather than copy number differences. |

In particular, MaAsLin 3’s feature-specific covariate testing directly implements our recommended best practices for differential expression testing in any metatranscriptomics analysis.^40^ To demonstrate this, we applied these improvements to bioBakery 3 profiles from the HMP2 IBDMDB metatranscriptomes.^25, 42^ Paired metatranscriptomic and metagenomic samples (383 CD samples from 52 individuals, 234 UC samples from 30 individuals, and 200 non-IBD samples from 27 individuals) were used to analyze expression of microbial pathways—coordinated biochemical reactions for a biological function (**Sup. Fig. 9**). In particular, pathway RNA relative abundances were regressed on the corresponding pathways’ DNA relative abundances (functional potential) in addition to disease status and dysbiosis, while controlling for age, antibiotic usage, read depth, and repeated sampling (**Methods**). Of the 209 significant (no model fitting errors, q-value *<* 0.1) diagnosis or dysbiosis associations identified, 142 (68%) were abundance associations, suggesting that while pathway expression varies in both abundance and prevalence, it predominantly differs in abundance (once controlled for functional potential).

To additionally demonstrate MaAsLin 3’s group and ordered covariate testing, all IBDMDB metagenomes from participants with Crohn’s disease (CD, *n* = 750) were used to regress species abundance and prevalence on food consumption frequency data. These were applied to both MetaPhlAn 4^43^ (recent) and MetaPhlAn 3^42^ profiles (as used in the IBDMDB publication; see **Methods**). Consistent with the group and ordered covariate models testing similar but distinct hypotheses in this setting, 59 of the 132 significant ordered associations had a corresponding group-wise association, and 59 of the 107 significant group-wise associations had a corresponding significant ordered association (MetaPhlAn 4). Of the 797 diet associations discovered in either MetaPhlAn 3 and MetaPhlAn 4, only 28 overlapped between the two, a result of differences in the abundance assigned to known taxa, differences in which taxa were included in each classifier, and differences in updated taxonomic naming. However, among the associations that did overlap, there was high agreement in the magnitude of the association (**Sup. Fig. 10**). One such association was the positive association between starch consumption in the last 4-7 days and *Eubacterium ventriosum* (SGB5045) prevalence, which is consistent with previous reports of reductions in other *Eubacterium* species during high protein, low carbohydrate diets that are depleted of starch.^44, 45^ Thus, MaAsLin 3’s linear model extensions expand the scope of inference in microbiome studies and identify plausible associations in diet data.

#### MaAsLin 3 clarifies and refines biomarker discovery in the inflammatory bowel diseases

To further explore associations in real data, we next applied MaAsLin 3 to the IBDMDB cohort to assess taxa associated with Crohn’s disease (CD) and ulcerative colitis (UC).^25^ Since the biomarkers of IBD are well established, we aimed to reconstruct generally accepted prior associations while assessing consistency across taxonomic classifiers and differential abundance testing methods. As above, MetaPhlAn 3 (as previously published)^25^ and 4 (updated) were used to construct taxonomic profiles (750 CD samples from 65 individuals, 459 UC samples from 38 individuals, and 428 non-IBD samples from 27 individuals), and the resulting profiles were tested for associations with disease status and dysbiosis while controlling for age, antibiotic usage, read depth, and repeated sampling (**Methods**).

Extending the previous analysis^25^ to account for differences in both the pediatric gut microbiome and IBD phenotypes, we stratified the IBDMDB cohort into populations that were pediatric (age *<* 16; 284 CD samples from 23 individuals, 95 UC from 8 individuals, and 161 non-IBD from 11 individuals) or adult (age *≥* 16; 448 CD samples from 37 individuals, 363 UC from 29 individuals, and 267 non-IBD from 16 individuals), with 16 corresponding to an age by which all participants had likely entered puberty.^46, 47^ The identified microbial biomarkers tended to show variable positive associations with IBD dysbiosis and diagnosis by age but more consistent negative associations. Of the 60 positive associations from either population (FDR q-value *<* 0.1, |*β*| *>* 1, and no model-fitting errors), only two overlapped (see **Methods** for criteria) between populations: *Citrobacter freundii* (SGB10083) and *Klebsiella pneumoniae* (SGB10115) with CD dysbiosis, both of which are thought to drive gut inflammation.^48, 49, 50^

Among the non-overlapping associations, 35 were unique to the adult population, with several microbes enriched in CD including several species of *Enterocloster* (*E. bolteae* (SGB4758), *E. aldenensis* (SGB4762), *E. lavalensis* (SGB4725), *E. clostridioformis* (SGB4760), and *E. citroniae* (SGB4761)) and several species from the class *Clostridia* (e.g. *Flavonifractor plautii* (SGB15132) and *Blautia faecicola* (SGB4867); **Fig. 4A**). Several well known IBD-associated microbes were positively associated with adult CD dysbiosis including *Enterococcus faecium* (SGB7967) and *E. faecalis* (SGB7962),^51,52, 53^ *Klebsiella oxytoca* (SGB10118),^54^ and *Hungatella hathewayi* (SGB4741), a microbe with substantial new links to inflammation.^55, 56^ Also consistent with previous studies, adult CD dysbiosis was associated with species from the genus *Clostridium* including *C. neonatale* (SGB6169), *C. perfringens* (SGB6191), *C. butyricum* (SGB6170), and two lesser characterized species (AT4 (SGB4753) and 1001270J-160509-D11 (SGB6139)).^51^ Fewer taxa were associated with UC (6) than CD (27), consistent with previous literature,^51^ but among UC associations, both *Blautia faecicola* (SGB4867) and *B. obeum* (SGB4811) were enriched.

**Figure 4:**
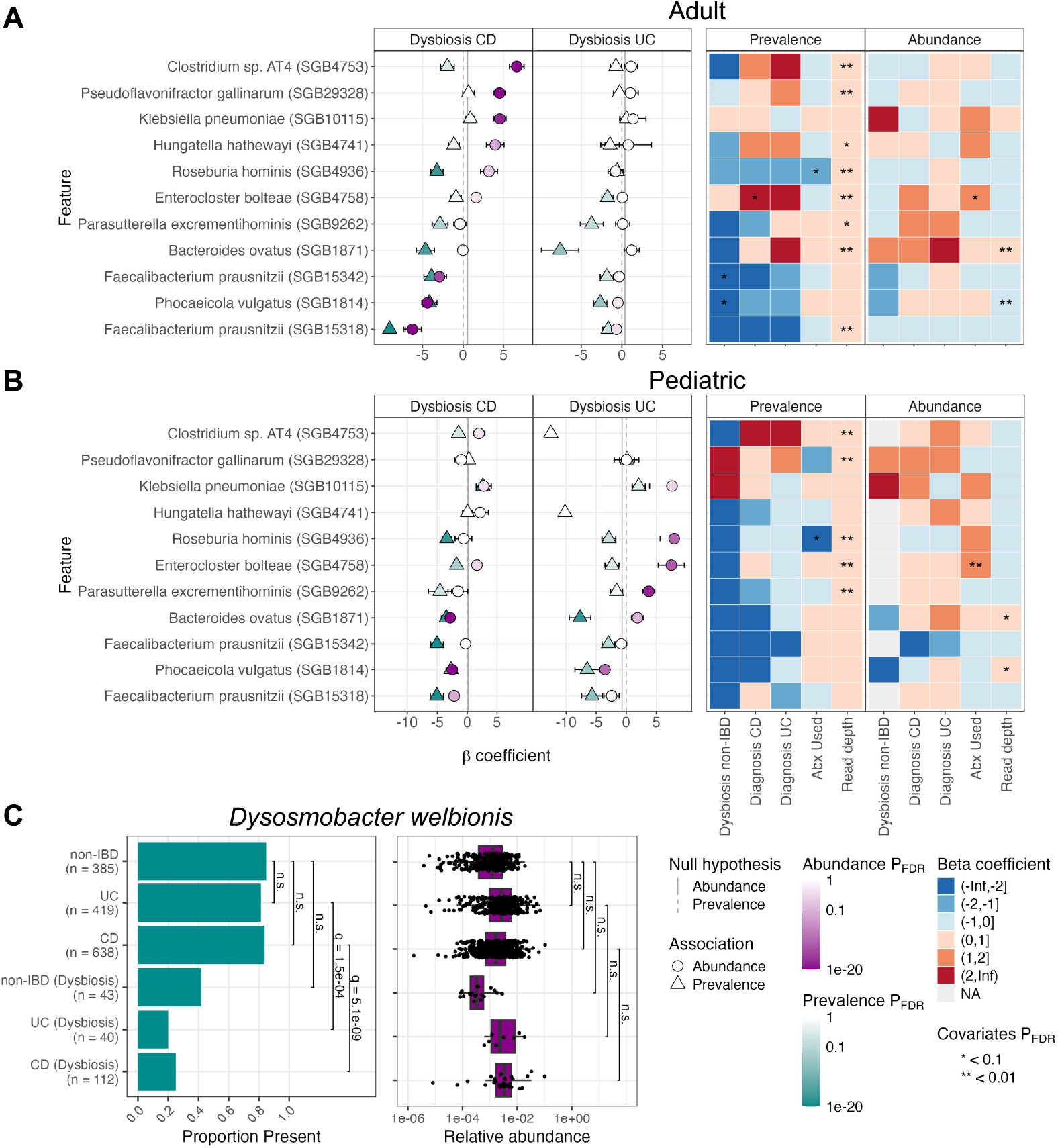
MaAsLin 3 applied to the HMP2 IBDMDB verifies and extends previous gut microbiome associations with IBD. The species-level abundances from the HMP2 cohort as determined by MetaPhlAn 4 were regressed in MaAsLin 3 using a model equivalent to that previously published^25^ incorporating disease-stratified dysbiosis, disease diagnosis, antibiotic usage, read depth, and a per-participant random intercept in individuals at least 16 years of age (**A**) or under 16 (**B**). Both panels show a default MaAsLin 3 output summary figure that has been subset to highlight species associated with either adult or pediatric dysbiosis. The estimated coefficients and their standard errors are represented by points and bars on the left. **C**. *Dysosmobacter welbionis* prevalence differed between dysbiosis and non-dysbiosis, while abundance did not differ. Bars show the comparisons evaluated in the MaAsLin 3 model after controlling for the aforementioned covariates and FDR correcting over all associations.

23 positive associations—microbes present or enriched during disease—were unique to the pediatric cohort including associations between CD dysbiosis and the previously implicated species *Escherichia coli* (SGB10068), *Ruminococcus gnavus* (SGB4584), *Bacteroides fragilis* (SGB1855 and SGB1853), and *Clostridium symbiosum* (SGB4699)^57, 58, 51^ (**Fig. 4B**). Interestingly, several species were enriched in pediatric CD dysbiosis that are typically found in probiotics including *Bifidobacterium breve* (SGB17247), *Bifidobacterium longum* (SGB17248), *Lactobacillus acidophilus* (SGB7044),^59, 60, 61^ along with a single SGB of *Faecalibacterium prausnitzii* (SGB15326, which was also associated with pediatric CD diagnosis).

By contrast with the small positive overlap, of the 223 negative associations that were significant in either adults or children, 53 (24%) of them overlapped including reductions in four *F. prausnitzii* species genome bins (SGB15316, SGB15318, SGB15332, SGB15342), *Phocaeicola vulgatus* (SGB1814), several *Bacteroides* species (*B. uniformis* (SGB1836), *B. ovatus* (SGB1871), *B. xylanisolvens* (SGB1867)), *Eubacterium rectale* (SGB4933), *Collinsella aerofaciens* (SGB14535), *Roseburia inulinivorans* (SGB4940), several *Alistipes* species (*A. shahii* (SGB2295), *A. putredinis* (SGB2318), *A. onderdonkii* (SGB2303)), *Blautia* species (*B. faecis* (SGB4820), *B. wexlerae* (SGB4837), and *B. massiliensis* (SGB4826)), and *Anaerostipes hadrus* (SGB4540). This confirms previous reports that typical gut commensals are lost during gastrointestinal inflammatory disorders.^51, 62, 63, 64^ Taken together, our findings confirm and expand our knowledge of IBD, differentiating both the gain and loss of specific microbes during inflammation, as well as the differences in these dysbioses among older and younger populations.

When MaAsLin 3 was run on the combined sample set from both age ranges, 372 significant associations were identified, of which 287 (77%) were prevalence associations. Of the 85 abundance associations, 50 were negative associations with either IBD diagnosis or gut dysbiosis, and, consistent with previous findings,^65, 66^ 89% (254) of the prevalence associations were negative. Since the sparsity was similar to the transcriptomic abundance data analyzed above (94.5% zeros in both), this suggests that in IBD, species themselves might predominantly differ in prevalence (e.g. gain or loss), while pathway expressions (once controlled for functional potential) predominantly differ in abundance.

As expected, more taxa were lost in CD (121) than in UC (80), though many overlapped including *Bacteroides uniformis* (SGB1836), *Eubacterium rectale* (SGB4933), and many species in the *Roseburia* genus. In particular, the reduction in the prevalence of *Dysosmobacter welbionis* (SGB15078) in CD and UC dysbiosis (**Fig. 4C**) highlights the synergy between MetaPhlAn 4’s taxonomic breadth and MaAsLin 3’s model capabilities. *D. welbionis* is a recently isolated human gut commensal that is associated with metabolic disorders in humans and can prevent diet-induced obesity in mice,^67, 68^ but it is newly detectable in MetaPhlAn 4 relative to MetaPhlAn 3. When analyzed with MaAsLin 2, *D. welbionis* showed a negative abundance association with both UC and CD dysbiosis, but when analyzed with MaAsLin 3, this association was shown to be driven only by a reduced prevalence of *D. welbionis* during dysbiosis, not by a reduced abundance when present (prevalence *β* = −4.14, −3.46; q-values = 1.5 *×* 10^*−*4^, 5.1 *×* 10^*−*9^; abundance *β* = −1.12, 0.15; q-values = 0.24, 1). Depending on the direction of causality, this may suggest that the organism’s phenotypic effects are exerted when it is present regardless of abundance, or that it is so sensitive to an inflamed gut that it is driven below the limit of detection during IBD.

When the other DA methods were applied to the MetaPhlAn 4 profiles, most associations identified by MaAsLin 3 overlapped with those of another method (83% were also identified by MaAsLin 2, 50% by ALDEx2, and 4% by ANCOM-BC2; **Sup. Fig. 11**). Among ALDEx2’s associations, 57% were unique to ALDEx2, since 203 of its 318 discovered IBD associations involved diagnosis, not dysbiosis (versus 35 of the 244 for MaAsLin 3). Among ALDEx2’s dysbiosis associations, 118 of the 127 were also discovered by MaAsLin 3. ANCOM-BC2 flagged only 11 associations as significant in the entire dataset, all of which overlapped with MaAsLin 3. These results support the plausibility of the new detail shed on the IBD gut microbiome by MaAsLin 3’s richer and more nuanced DA model components.

## Discussion

Differential abundance testing is a key component in most microbial community analyses, but ambiguity remains regarding the best approaches.^1^ In large part, this is driven by the challenging properties of microbiome data, particularly compositionality, sparsity, and high dimensionality. To address these, we introduced MaAsLin 3 to discover differential abundance and prevalence associations while accounting for compositionality, experimental protocols determining absolute abundance, and new covariate types (including metatranscriptomics). On simulated datasets, MaAsLin 3 outperformed current state-of-the-art DA methods, maintaining precision as well as or better than other methods, even when its modeling assumptions were violated, and achieving higher recall in most simulations than any method with similar precision. Additionally, when estimating absolute abundance coefficients from relative abundance data, MaAsLin 3 and ANCOM-BC2 produced coefficients that were the most accurate (**Fig. 1C, Sup. Fig. 2**). However, only MaAsLin 3 includes the ability to natively handle absolute abundance protocols (both spike-in and total biomass quantification) that are required to determine these coefficients experimentally. When applied to the HMP2 IBDMDB population of participants with and without IBD, 77% of taxonomic associations with IBD were quantified as prevalence associations, and 44 associations were new to MaAsLin 3 compared to MaAsLin 2, eight of which were not previously significant due to opposing abundance and prevalence associations (**Fig. 4**).

MaAsLin 3’s most prominent methodological improvements include median adjustment for compositional relative abundances and separate models for prevalence (presence/absence) and abundance associations. The former provides a simple way to improve on MaAsLin 2’s minimal log transform that is equivalent to experimental absolute abundance data given reasonable assumptions, thus providing clear benefits. The latter can be beneficial both statistically and biologically. Statistically, modeling prevalence can help avoid the pitfalls of zero imputation for sparsity.^16, 28, 29^ Furthermore, the predominance of prevalence associations in real data suggests that many taxa have abundances that are low enough or variable enough that only associations involving complete loss of the taxon or a reduction below the detection threshold are strong enough to be observed. Biologically, microbes with unique niches can exert phenotypic effects when simply present even at very low relative abundances, such as methanogenesis unique to the archaea^69^ or the ability of *Clostridiodes difficile* to cause infections even in very low doses.^19^ Further, prevalence associations typically agreed with their corresponding abundance associations, so they might provide another measure of a taxon’s general ability to survive in a particular environment.

Relative to previous studies of this cohort, more precise taxonomic characterization using MetaPhlAn 4 allowed us to further elucidate biomarkers of IBD. The discovery of a prevalence-only association between *D. welbionis* and IBD dysbiosis highlights the specificity gained by splitting abundance from prevalence associations (**Fig. 4C**). *D. welbionis* is a recently isolated human commensal; it has been hypothesized to have a beneficial role in metabolic disorders based on observational human studies and on decreased weight gain during a high fat diet in mouse models.^67, 68^ Of interest in IBD, some CD is characterized by creeping fat,^70, 71^ the etiology of which is unknown, but since *D. welbionis* has been implicated in reducing white adipose fat hypertrophy,^67^ the presence of this microbe in non-dysbiotic CD individuals might reduce the risk of creeping fat. Additionally, a recent study found that *D. welbionis* metabolizes cholesterol and possesses a protein similar to IsmA, an enzyme that converts cholesterol to coprostanol and is strongly associated with depleted cholesterol and increased coprostanol in human stool samples.^72^ Thus, the loss of *D. welbionis* during dysbiosis might reduce the microbial community’s ability to metabolize cholesterol into coprostanol, a previously reported clinical outcome in IBD.^73, 74^ This may also suggest a mechanistic link by which the gut microbiome could mediate some of the increased risk of cardiovascular disease in IBD patients.^75^

We also sought to determine whether features of the gut ecosystem might link to the observation that the severity and progression of IBD tends to be worse in pediatric patients.^76^ As in most early onset chronic diseases, this is a setting in which causality can be difficult to determine: do gut microbiome changes drive this increased severity, or does more severe disease tend to manifest earlier in life? Perhaps the most relevant finding was that microbes typically hypothesized to drive inflammation were more likely to be enriched in the gut microbiome of adolescents. If gut microbial communities remain more plastic in adolescents than adults,^47, 77^ this adolescent-associated instability might allow invasion of opportunistic, fast-growing microbes more readily than in adults,^78^ thus forming a cause of more severe disease. This effect might be exacerbated by low fiber diets, a hallmark of children in the US,^79^ since recent studies have suggested that a shift from microbes utilizing carbohydrate to mucin was able to stimulate IBD phenotypes.^80^ This combination of early life gut instability, opportunistic invasion, and suboptimal diet could also have lasting impacts on the hosts’ immune system.^81^ While the severity and faster progression of childhood IBD is likely multifaceted, these models suggest that gut instability and the invasion of IBD-typical microbes might play a more important role in pediatric IBD.

In addition to improving the core features of compositionality, abundance, and prevalence detection, MaAsLin 3 supports a variety of new inference types designed to meet the needs of the diverse research conducted on microbial communities. Group difference and level-versus-level difference tests address the same scenarios as ANCOM-BC2^12^ and can be applied to testing differences across dietary groups, disease classifications, or collection sites. Contrast tests are useful for testing all pairwise differences between groups such as in analysis of IBD subtypes.^82^ Finally, feature-specific covariates are particularly useful for controlling for per-gene DNA abundance when regressing per-gene RNA abundance in metatranscriptomic studies.^83, 40^ These expanded capabilities allow MaAsLin 3 to accommodate complex experimental designs, without the need for them to be computationally coded in a complex manner.

Despite these improvements, some limitations still remain. While MaAsLin 3 outperformed previous methods on essentially all simulations considered here, there remain other plausible microbial data distributions not yet tested. Additionally, many DA methods exist beyond those analyzed here; for brevity, we only compared against some of the most commonly used.^10, 13,11^ Notably, all DA methods remain affected by lower recall in small samples and by the necessary limitations of finite read depths. Compared to MaAsLin 2 and ANCOM-BC2, MaAsLin 3 achieved lower recall on small sample sizes because logistic models often require more samples to reveal a significant association than linear models,^84, 85^ and its linear modeling portion only uses the non-zero subset of the data (**Sup. Fig. 2**). However, the slightly higher recall of these other methods in low sample sizes came at the cost of more biased coefficient estimates (MaAsLin 2) and severely reduced precision (ANCOM-BC2). Second, limited read depth prevented MaAsLin 3 from always distinguishing correctly between abundance and prevalence associations and sometimes caused it to miss associations. When a rare feature’s abundance is associated with a covariate, that feature might be more likely to drop below the limit of detection depending on the covariate’s value, yielding an entirely missed effect or a spurious prevalence effect in place of a true abundance effect. Rigorously accounting for this phenomenon would require experimental techniques such as culture-enriched molecular profiling,^86^ but in their absence, MaAsLin 3 reports prevalence associations deemed likely to be spurious and provides diagnostic plots for manual curation when necessary.

In summary, the methods introduced in MaAsLin 3 represent an important advancement in the accuracy, complexity, and biological detail achievable in microbiome DA testing. They out-perform state-of-the-art DA methods across a variety of simulations when testing for traditional associations, and MaAsLin 3 expands the space of possible associations with a variety of new covariate types. When applied to real datasets, MaAsLin 3 discovered hundreds of biologically plausible and internally consistent microbial associations when applied to adult and pediatric IBD. Many of these aligned with previous IBD findings, but some were only detectable with these updated models, which also clarified that most gut microbiome changes during inflammation corresponded with complete microbial loss. This informs, for example, strategies such as live biotherapeutic microbial supplementation in favor of e.g. prebiotic or dietary management in such settings.^87^ MaAsLin 3 thus provides researchers with an important and much-improved set of capabilities for understanding microbiome associations with environmental phenotypes, human health, and disease.

## Data Availability

MaAsLin 3 is available as an R package at https://github.com/biobakery/MaAsLin3, and it is in submission at Bioconductor. All scripts used in benchmarking and real data analysis are available at https://github.com/willnickols/MaAsLin3_benchmark. Tutorials for MaAsLin 3 are available at https://github.com/biobakery/biobakery/wiki/MaAsLin3/ and https://github.com/biobakery/biobakery/wiki/MTX-model-3.

## Acknowledgments

The computations in this paper were run in part on the FASRC Cannon cluster supported by the FAS Division of Science Research Computing Group at Harvard University. The work was supported by the National Institute of Diabetes and Digestive and Kidney Diseases of the National Institutes of Health (R24DK110499) to C.H. and the National Institute of Allergy and Infectious Diseases (U19AI110820) to D. Rasko (to C.H.) and the Crohn’s and Colitis Foundation Early Career grant (K.N.T).

## Methods

The MaAsLin 3 algorithm consists of (1) total-sum scaling feature abundance profiles to obtain relative abundances, (2) generating a prevalence (present versus absent) profile and retaining separately the non-zero abundances, (3) log transforming (base 2) the abundance dataset, (4) performing an augmented logistic regression on the prevalence dataset and linear regression on the abundance dataset, and (5) combining the logistic and linear effects into an overall effect for each feature-metadatum pair. This is in contrast to MaAsLin 2, which total-sum scales the feature abundance profiles, sets zeros to half the minimum observed abundance, log transforms all abundances (including imputed values), and performs a linear regression on the transformed values.

### MaAsLin 3 workflow

MaAsLin 3 requires, at a minimum, a samples-by-features table of feature abundances (counts or pre-computed abundances for taxa, genes, etc.) and a samples-by-covariates table of metadata. The table of features is then normalized (by default, total-sum scaling to produce relative abundances) and features are optionally filtered by thresholds for prevalence, abundance, and minimum variance for non-zero abundances. By default, any feature present with non-identical abundance in at least two samples is retained. The abundance table is then split into a presence/absence table and a table of non-zero values that are subsequently transformed (log base 2 transformed by default). The specified logistic and linear models are fit on the presence/absence mask and transformed data respectively, and the results are saved. A summary plot is produced for the top associations, and association-specific diagnostic plots are created for the top associations.

#### Abundance modeling

For the abundance associations, a linear model (possibly with mixed effects) is fit based on the specified formula one feature at a time. Following the notation of ANCOM-BC, let *i ∈* {1, …, *m*} be the feature index, *j ∈* {1, …, *g*} be the covariate index, and *k ∈* 1, …, *n* be the sample index. Let 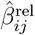 be the slope corresponding to covariate *j* when regressing the transformed relative abundance of feature *i*. When testing relative abundance associations with the median correction off, the 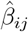 are each tested against a null of zero according to standard homoskedastic regression theory. If the median comparison is on for testing absolute abundance associations (as by default), the median of the coefficients for each covariate 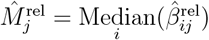 is calculated, and the coefficients 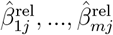 are tested against 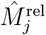 with a test that accounts for the variability in both the coefficients and the median (**Supplementary Information**). Optionally, this median 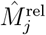 is subtracted from each 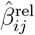.

In its normalization step, MaAsLin 3 can incorporate absolute abundance data from both spike-in experiments and total abundance estimation procedures. Let *A*_*ik*_ be the absolute abundance of feature *i* in sample *k*, let *T*_*k*_ = Σ_*i*_ *A*_*ik*_ be the total absolute abundance in sample *k*, and let *P*_*ik*_ = *A*_*ik*_*/T*_*k*_ be the relative abundance of feature *i* in sample *k*. Note that the ratios of relative abundances are equal to the ratios of absolute abundances:^88^ *P*_*ik*_*/P*_*i*_*′*_*k*_ = *A*_*ik*_*/A*_*i*_*′*_*k*_. In a spike-in experiment, estimators of both the absolute abundance 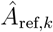 and the relative abundance 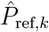 for the spike-in reference feature are known. Therefore, MaAsLin 3 can take as input the vector of absolute abundances of the spike-ins 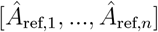 and use these to estimate the absolute abundances of the non-reference features as 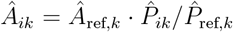. Alternatively, MaAsLin 3 can take estimates 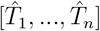 of the samples’ total abundances and estimate the absolute abundances as 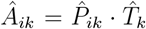. These absolute abundances are then filtered, log (base 2) transformed, and regressed as in the relative abundance case but without the median comparison. As described previously,^89^ the spike-in procedure will eliminate bias due to sampling efficiency while the total abundance scaling procedure will not.

Let 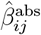 be the slope corresponding to covariate *j* when regressing the transformed absolute abundance of feature *i*. As described in the **Supplementary Information**, when there is no sparsity, the absolute abundance slope can be decomposed as 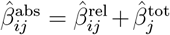 where 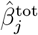 is the slope corresponding to covariate *j* in a regression of the log (base 2) transformed persample total abundances on the same covariates. Since 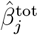 does not depend on the feature *i*, the difference in two features’ relative abundance coefficients is equal to the difference in the features’ absolute abundance coefficients, and the orders of the slopes over the features are the same. Furthermore, as described in the **Supplementary Information**, if at least half the features are not changing on the absolute scale 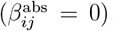, the test of 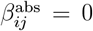 for any individual coefficient is equivalent to a test of 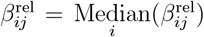, hence the test of the relative abundance coefficients against their median. Equivalent results hold even with unequal sampling efficiencies (**Supplementary Information**). These results have assumed no sparsity, but equivalent results can be obtained with sparsity if alternative, typically weaker, assumptions are satisfied (**Supplementary Information**).

#### Prevalence modeling

For the prevalence associations, a logistic model or logistic model with mixed effects is fit based on the specified formula one feature at a time. By default, a data augmentation scheme is performed to avoid linear separability and outsized influence from a small number of data points. Let **B**_*i*_ be the *n×*1 binary vector of presence/absence values for feature *i*. Let **X** be the *n×* (*g* + 1) fixed effects design matrix from *g* covariates and an intercept. The data augmentation procedure creates a 3*n×*1 augmented presence/absence vector 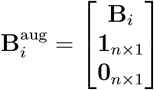 and a 3*n* × (*g* + 1) augmented design matrix 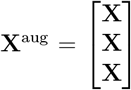 and performs weighted logistic regressions with weights of 1 for the first *n* (original) values and weights of *g/*(2*n*) for the remaining (augmented) values. In this way, each original value is augmented with an extra presence and an extra absence such that the total weight of the augmented samples is equivalent to *g* additional samples. This technique, equivalent to the Diaconis-Ylvisaker prior^34, 35^ has been proposed as an alternative to Firth regression with the advantage that it extends easily to mixed effects models and model comparisons procedures.

When a low-abundance feature’s abundance is associated with a metadatum, a spurious prevalence association can be induced by the abundance association if the feature regularly falls below the limit of detection despite being non-zero. The exact relationship between an abundance association and the induced prevalence association is complex, depending on both the (unknown) abundance association and the read depth. While a full treatment of this problem is beyond the scope of this paper, the following heuristic was implemented. After the prevalence associations are fit, a linear regression is performed on the log-transformed relative abundances, and the coefficients are tested against zero (not the median). Then, each prevalence association is flagged as likely abundance-induced if (1) the corresponding abundance association has a significant q-value, (2) the sign of the abundance coefficient is the same as the sign of the prevalence coefficient, and (3) the magnitude of the abundance coefficient is larger than the magnitude of the prevalence coefficient.

Typically, read depth should be included as an untransformed covariate. Assuming the distribution of reads for a feature with relative abundance *r* in a sample with *n* reads is approximately Binom(*n, r*), the prob(ability of the feature being sampled is 1 *−* (1 *− r*)^*n*^, so the log odds of inclusion are: log 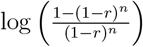. For *r* small and *n* large (e.g., *n >* 2*/r*), 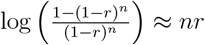, so the log odds of a feature being present are approximately linear with the read depth, motivating the inclusion of the untransformed read depth covariate. When the read depth is independent of the other covariates (as in the simulations), it should not be included because of the non-collapsibility of the odds ratio.^90^ Otherwise, if it is associated with the other covariates as is often the case due to differences in sample biomass and extraction, it should be included.

#### Combining associations

To test for an overall association between a feature and a covariate, particularly for comparing against other tools that do not distinguish abundance from prevalence, an overall p-value is calculated as the Beta(1, 2) cumulative density function evaluated at the minimum of the two p-values from the abundance and prevalence models. Under the null of no abundance or prevalence association between the feature and the covariates, both models produce p-values uniform on 0 to 1.^91^ Furthermore, these p-values are independent since only the abundances in samples with the feature present are used to determine the abundance p-value, and knowing the feature is present (the only information used in the logistic p-value) gives no information about what its non-zero abundance is. Since the minimum of two independent Unif(0, 1) random variables has a Beta(1, 2) distribution, the minimum of the two p-values will have a Beta(1, 2) distribution under the null.

#### New model components and specification capabilities

MaAsLin 3 allows general mixed effects models through the formula parsing of the package lme4,^41^ including interaction terms and random intercepts.

For omnibus testing of whether all coefficients corresponding to categories of a covariate are zero, an ANOVA-style test is performed. For linear models, this is an F-test (performed with the package lmerTest^92^ for mixed effects models), and for logistic models this is a likelihood ratio test.

For testing differences in levels of an ordered covariate, contrast tests are performed between consecutive levels in the fit model with a right hand side corresponding to zero or the median coefficient difference between the relevant levels. This test is performed with the package lmerTest for linear mixed effects models and with the package multcomp^93^ otherwise.

Contrast tests are performed in the same way as the tests for ordered covariates but with arbitrary user-specified contrast matrices.

A feature-specific covariate table can be provided with dimensions matching the abundance table. Then, in the regression for feature *i*, the *i*^*th*^ column of the table will be extracted and used as a covariate in the expanded *n ×* (*g* + 2) design matrix **X**.

### Benchmarking on simulated microbial community data

#### SparseDOSSA 2

The R package SparseDOSSA 2 was used to parametrically generate synthetic microbial community abundances according to templates informed by real data.^5^ Using the “Stool” template based on real microbiome data,^94^ SparseDOSSA 2 generated underlying null distributions of features from zero-inflated log-normal distributions. Then, synthetic metadata was created by sampling from a multivariate normal distribution and converting half the covariates to binary values. Using this synthetic metadata along with a synthetically generated table of associations to add in (effect sizes uniform on 2.5 to 5 unless otherwise specified), abundance (log-linear) and prevalence (logistic) associations were imposed on the absolute scale, and reads were sampled from the resulting relative abundances.

To evaluate MaAsLin 3’s performance on group and ordered predictors, a multi-level categorical covariate was thermometer encoded in the synthetic metadata matrix (i.e. for an ordered covariate with levels 1 through *p* and an observed level of *j, p* indicators are included with the first *j* set to 1 and the rest set to 0). Next, coefficients were generated representing the total difference between the baseline level and the level with the greatest difference, and a Dirichlet(**1**) draw was used to divide the total difference among the levels. Then, with probability 1/2, each level’s coefficient was set to 0 and added to the subsequent coefficient. Finally, the resulting metadata and coefficients were used to generate synthetic abundances, and the thermometer columns were collapsed into a single ordered categorical covariate.

Natively, SparseDOSSA 2 does not simulate spike-ins. Therefore, SparseDOSSA 2 was modified to include an extra feature in the absolute abundance generation step with an abundance chosen uniformly from 1% to 10% of the total abundance. The absolute abundance of this spiked-in feature was then stored and used in MaAsLin 3’s spike-in mode. After this feature was added on the absolute scale, read sampling was performed as before.

#### ANCOM-BC generator

The data generation strategy from ANCOM-BC’s benchmarking was also applied with minor modifications.^11^ Using a template from a previously sequenced soil community as the null distribution,^95^ two groups of samples were created with the mean absolute abundances in one group multiplied by effect sizes drawn uniformly between 2.5 to 5 to create an unbalanced microbial load. Then, structural zeros were added in by setting a random 20% of features to 0 for all samples in a group. Finally, reads were sampled from the resulting profiles using rarefaction subsampling. This sampling used mock sequencing depths with the same distribution as in the ANCOM-BC evaluation but scaled to a mean depth of 50,000 reads for consistency with the other simulations.

#### Running differential abundance tools on simulated datasets

For simulations in which MaAsLin 3 was run on only relative abundance data, MaAsLin 3 (version 3.0.0) was almost always run with all default settings (median comparison on, data augmentation on, warning on abundance induced prevalence, no spike-ins) and a formula incorporating all simulated covariates (i.e. a correctly specified model). The first exception was that for repeated sampling scenarios, per-subject random intercepts were also used. The second exception was that, in all relative abundance simulations, median subtraction was turned on (i.e. subtracting the coefficient medians after testing against them) to evaluate the bias of the resulting differences as estimators of the synthetic absolute scale coefficients. (By default in the software, this is off so that the relative abundance coefficients are returned for ease of interpretation.) When run using absolute abundance spike-in information, MaAsLin 3 was run with the spike-in absolute abundance provided as a parameter and the median comparison off.

ALDEx2 (version 1.34.0) was run with a model incorporating all the covariates (a correctly specified model) and default settings. Since ALDEx2 does not provide a random effects option, to control for correlated measurements in simulations involving repeated sampling, per-subject fixed effects were fit and then removed from subsequent analysis (not included in the FDR correction etc.). P-values were FDR corrected with the Benjamini-Hochberg procedure.^96^

ANCOM-BC2 (version 2.4.0) was run with a model incorporating all the covariates (a correctly specified model) and default setting except for (1) an FDR level of 0.1 rather than 0.05, (2) a prevalence threshold set to 0 rather than 0.1, and (3) the Benjamini-Hochberg FDR procedure rather than the Holm procedure, all for consistency with the other tools. Because it produced many false positives, the structural zero option was left off except in the ANCOM-BC generator evaluation when the assumptions for this option were satisfied. Only significant associations that passed ANCOM-BC2’s pseudocount sensitivity screen were counted as significant. For repeated sampling scenarios, per-subject random intercepts were included. Coefficients were converted from their default log_*e*_ scale to the log_2_ scale for consistency with other tools in the analysis.

MaAsLin 2 (version 1.16.0) was run with a model incorporating all the covariates and default settings except for (1) an FDR level of 0.1 rather than 0.25 and (2) abundance and prevalence thresholds set to 0, both for consistency with the other tools. For repeated sampling scenarios, per-subject random intercepts were used.

### Benchmarking on real data

#### Randomization procedure

The binary metadata labels of the 38 datasets^1^ were randomly shuffled to create 100 mock datasets for each real dataset. MaAsLin 3 was then run on both the randomized datasets and the original (non-randomized) datasets.

#### Real absolute abundance data

Three datasets with inferred absolute abundance data were analyzed: an infant gut dataset that used a spike-in procedure,^36^ a mouse diet dataset that used 16S digital PCR to estimate total microbial load,^37^ and an IBD/PSC dataset that used flow cytometry to estimate total microbial load.^38^ For all three of these studies, estimated per-sample, per-taxon absolute abundance tables were already available, so the datasets were analyzed without further normalization in MaAsLin 3 to obtain absolute abundance coefficients. That is, equivalent results could have been obtained by supplying the spike-in abundances or total microbial load estimates to MaAsLin 3 along with the relative abundance tables, but since the absolute abundance scaling operations MaAsLin 3 would have performed were already precomputed, the data were used directly. For the infant dataset, the model included days since birth and read depth as fixed effects and infant ID as a random effect. For the mouse diet dataset, the model included diet and day as fixed effects and mouse ID as a random effect (read depth was not included because all samples had equal read depths). ANCOM-BC2 produced errors when the mouse ID was included as a random effect, so it was removed from the model. (This removal seemed to have a minimal effect since ANCOM-BC2’s coefficients still achieved essentially perfect accuracy in this setting). For the IBD/PSC dataset, diagnosis, age, gender, BMI, and read depth were included as fixed effects with diagnosis as a categorical variable that compared PSC-only, PSC-UC, CD-only, and PSC-CD against healthy controls as the baseline. Read depth was included as a covariate in the real datasets because deeper sequencing will often associate with higher prevalence (taxa are detected more often with more reads), and this deeper sequencing could be confounded with the key sample covariates.

#### HMP2-IBDMDB analysis

The HMP2 Inflammatory Bowel Disease Multi-omics (IBDMDB) dataset derives from 130 individuals recruited in five US medical centers with CD, UC, or no inflammatory bowel disease, who donated stool samples for one year, up to 24 times each.^25^ Raw sequencing data from the 1637 metagenomic samples were cleaned of host contaminant reads and low-quality reads with KneadData^42^ using default parameters. Next, MetaPhlAn 4.0.6 (Oct22 database)^43^ was used to construct taxonomic profiles with species genome bins for increased coverage, and the MetaPhlAn 3.0 profiles were downloaded from IBDMDB.^25^ The resulting abundances were analyzed with each tool with antibiotic usage, diagnosis, and dysbiosis status as fixed effects and participant ID as a random effect. Dysbiosis status is intended to reflect IBD disease flares and was calculated as previously described^25^ but with all controls classified as non-dysbiotic. When not already split by pediatric and adult populations, age was also included as a fixed effect. Based on previous benchmarking presented in this manuscript, associations were only considered significant if they had a q-value below 0.1, a coefficient with an absolute value greater than 1, and no model fitting errors. For each feature-metadatum pair, the association was considered to overlap between the adult and pediatric populations if the association’s more significant coefficient (abundance or prevalence) from each population had the same sign.

Using the subset of samples from participants with CD, abundances were regressed in MaAsLin 3 using a model that incorporated categorical dietary frequency information (dietary component eaten in the last 7 days, in the last 4-7 days, etc.) as a group or ordered predictor. Also included in this model were dysbiosis, antibiotic usage, age, and read depth as fixed effects and participant ID as a random intercept.

Previously computed pathway abundances from metagenomic and metatranscriptomic data using HUMAnN 3^42^ were downloaded from IBDMDB.^25^ The abundances were pre-processed in MaAsLin 3 by, for each pathway in each sample, (1) using the relative abundances as-is if the DNA abundance was non-zero, (2) using the RNA abundance as-is and setting the DNA abundance to the log_2_ of the minimum non-zero DNA relative abundance in the dataset divided by 2 if the RNA was non-zero but the DNA was zero, or (3) excluding the observation if both the DNA and RNA were zero. Case (1) matches the usual assumptions of MaAsLin 3; case (2) assumes that the pathway DNA abundance was below the limit of detection; and case (3) assumes the pathway was not present at all, so there is no expression information. Using MaAsLin 3, the metatranscriptomic relative abundances were regressed on age, antibiotic usage, diagnosis, and dysbiosis status as fixed effects, participant ID as a random effect, and pathway DNA relative abundance as a covariate-specific fixed effect, following previous recommendations.^40^ Only relative abundance associations were of interest for the metatranscriptomics analysis, so no median comparison was performed.

## Supplementary Information

### Relative and absolute coefficients differ by a constant shift when extraction efficiency is equal and all features are present

First, consider the case in which all features have equal extraction efficiency and there is no sparsity. Following the notation of ANCOM-BC, let *i ∈ {*1, …, *m}* be the feature index, *j ∈ {*1, …, *g}* be the covariate index, and *k ∈ {*1, …, *n}* be the sample index. Let **X** be the *n×* (*g* +1) design matrix of the metadata (including the intercept). With *A*_*ik*_ as the absolute abundance of feature *i* in sample *k*, let *Z*_*ik*_ be the log_2_ absolute abundance, so *Z*_*ik*_ = log_2_(*A*_*ik*_). Also, suppose the absolute abundance is related to the metadata by: *E*(*Z*_*ik*_) = Σ_*j*_ *X*_*kj*_*β*_*ij*_ +*ϵ*_*ik*_ where *β*_*ij*_ is the slope relating covariate *j* to feature *i*’s log_2_ absolute abundance and *ϵ*_*ik*_ is some error with mean 0. These slopes can be grouped together for a feature as a (*g* + 1) *×* 1 column vector: ***β***_*i*_ = (*β*_*i*0_, *β*_*i*2_, …, *β*_*ig*_)^*T*^. Let the *n ×* 1 vector **Z**_*i*_ = (*Z*_*i*1_, *Z*_*i*2_, …, *Z*_*in*_)^*T*^ denote the vector of log_2_ absolute abundances for feature *i*. With *T*_*k*_ = Σ*i A*_*ik*_ as the total absolute abundance in sample *k*, let *D*_*k*_ = log_2_(*T*_*k*_). With *P*_*ik*_ = *A*_*ik*_*/T*_*k*_ as the relative abundance of feature *i* in sample *k*, let *Y*_*ik*_ = log_2_(*P*_*ik*_), so 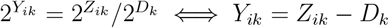. As with the absolute abundances, let **Y**_*i*_ be the *n ×* 1 vector of log_2_ relative abundances for feature *i*, and let **D** be the *n ×* 1 vector of log_2_ total abundances. Thus, **Y**_*i*_ = **Z**_*i*_ *−* **D** for all *i*.

When observing a vector **Z**_*i*_ and wanting to estimate ***β***_*i*_, the ordinary least squares method is typically used: 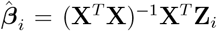. By the abundance decomposition above, this also gives:

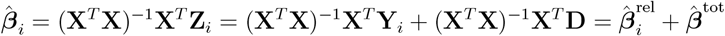

where 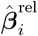 is the result of regressing the log_2_ relative abundances on the design matrix, and 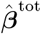 is the result of regressing the log_2_ total abundances on the design matrix. Note that since 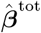 is the same for all features, if the absolute abundance coefficient for one feature *i* is *d* larger than the absolute abundance coefficient for another feature *i*^*′*^ (i.e., 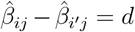), the relative abundance coefficient for feature *i* will be *d* larger than the relative abundance coefficient for feature *i*^*′*^ (i.e., 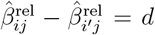). Thus, if all that is available is the relative abundance data, absolute slopes 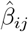 themselves cannot be determined. However, the relative coefficients 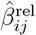 *can* be determined, and the ordering of and spacing between these coefficients will be identical to the ordering of and spacing between the absolute coefficient for each metadatum.

Since the OLS solution is unbiased for ***β***_*i*_, the expectations will be equal too:

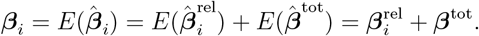

Assuming at least half of the features do not change with respect to a particular metadatum (i.e., *β*_*ij*_ = 0 for at least half the features *i*), the median absolute abundance coefficient will be 0 (i.e., med(*β*_1*j*_, *β*_2*j*_, …, *β*_*mj*_) = 0), as has been previously noted in LOCOM.^31^ Expanding each of these coefficients using the expectation decomposition above gives

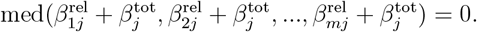

Since the term 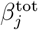 is the same in all of these, this implies 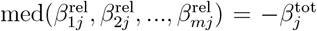. Thus, a test of *β*_*ij*_ = 0 is algebraically equivalent to the tests:

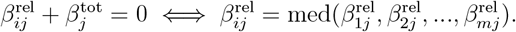

That is, testing whether one feature’s relative abundance slope for a metadatum is different from the median relative abundance slope for that metadatum is the same as testing whether that feature’s absolute abundance slope is different from 0. This motivates the median comparison test implemented in MaAsLin 3.

### Covariate slopes are unaffected by extraction efficiency

Now, let the extraction efficiency differ by feature, but assume each feature’s extraction efficiency depends only on the feature, not on what else is in the sample. Let *A*_*ik*_ and *Z*_*ik*_ be the true absolute and log absolute abundances as above, and let *E*_*i*_ be the sampling efficien cy of feature *i* with *S*_*i*_ = log_2_(*E*_*i*_). Now, let *T*_*k*_ = Σ_*i*_ *A*_*ik*_*E*_*i*_ and *D*_*k*_ = log_2_(*T*_*k*_), so 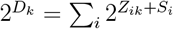. Then, the relative abundances can be written as.

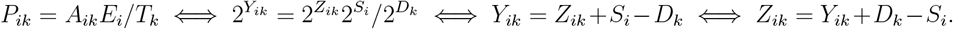

Let **S**_*i*_ be *S*_*i*_ repeated *p* times to match the dimensions of the feature-specific slopes. With an equivalent decomposition to before,

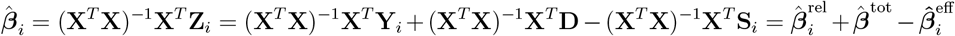

where 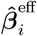 is the regression of the log_2_ sampling efficiency on the covariates **X**. However, since all the elements of **S**_*i*_ are equal, 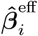 will just be an intercept and then zeros for all the slopes corresponding to covariates (i.e., 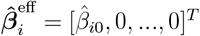). Thus, excluding the intercept, the same result as before holds: 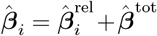. Therefore, the same results as above follow, justifying the median comparison technique and its equivalence to absolute testing when at least half the features are unassociated with a metadatum.

### With sparsity

For simplicity, consider the equal extraction efficiency case again. Before, the coefficients could be decomposed as 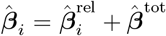 where 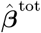 was the same for all features, but this was reliant on the data matrix **X** being the same for all features. When some features have zero abundances and only non-zero abundances are included in the regression, there will be different matrices **X**^(*i*)^ for each feature. Thus, the equation relating the slopes is 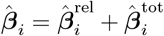 with an index on the total abundance regression coefficient. Since 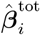 can now differ by feature, the ordering and spacing of the 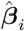 are not necessarily the same as of the 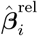. To proceed, note that the following quantities are being estimated with these regressions:

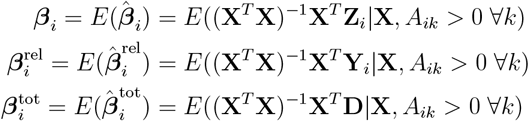

Since 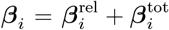, we would retain the consistent ordering and spacing properties, at least in expectation, if

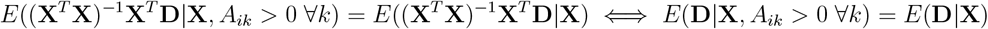

for all *i*. That is, the results before would hold if (1) all features are always present (no sparsity) or (2) the expected total abundance is independent of whether any particular feature is actually present (i.e., the conditional expectation of **D** is the same with or without the conditioning on *A*_*ik*_ *>* 0 ∀*k*). When the carrying capacity of a community is determined by the covariates and the community is at carrying capacity (e.g., in an adult gut), this is more likely to be true. However, during periods of colonization (e.g., in the infant gut), this is unlikely to hold.

### Test implementation

The test statistic is 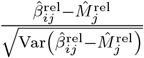. The variance can be decomposed: 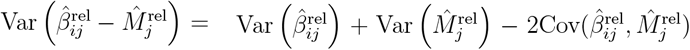. First, 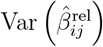 is already estimated in the model fitting. Second, 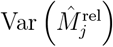 is estimated as 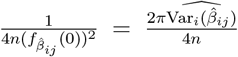 based on the asymptotic distribution of the median assuming the 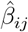 are i.i.d. from a normal distribution with mean 0.^91^ While this will almost always be violated practically, the estimate of the 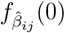 will typically be too small (the estimated density will be too dispersed), resulting in a conservative rather than anti-conservative test. Third, 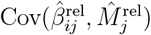 is estimated by bootstrapping the 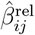 from their approximate normal distributions and computing 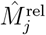. The computed test statistic is then evaluated against the distribution of 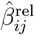 to determine a p-value. While the distribution will not be exact, it will typically be close to a standard normal.

## Supplementary Figures

**Supplementary Figure 1:**
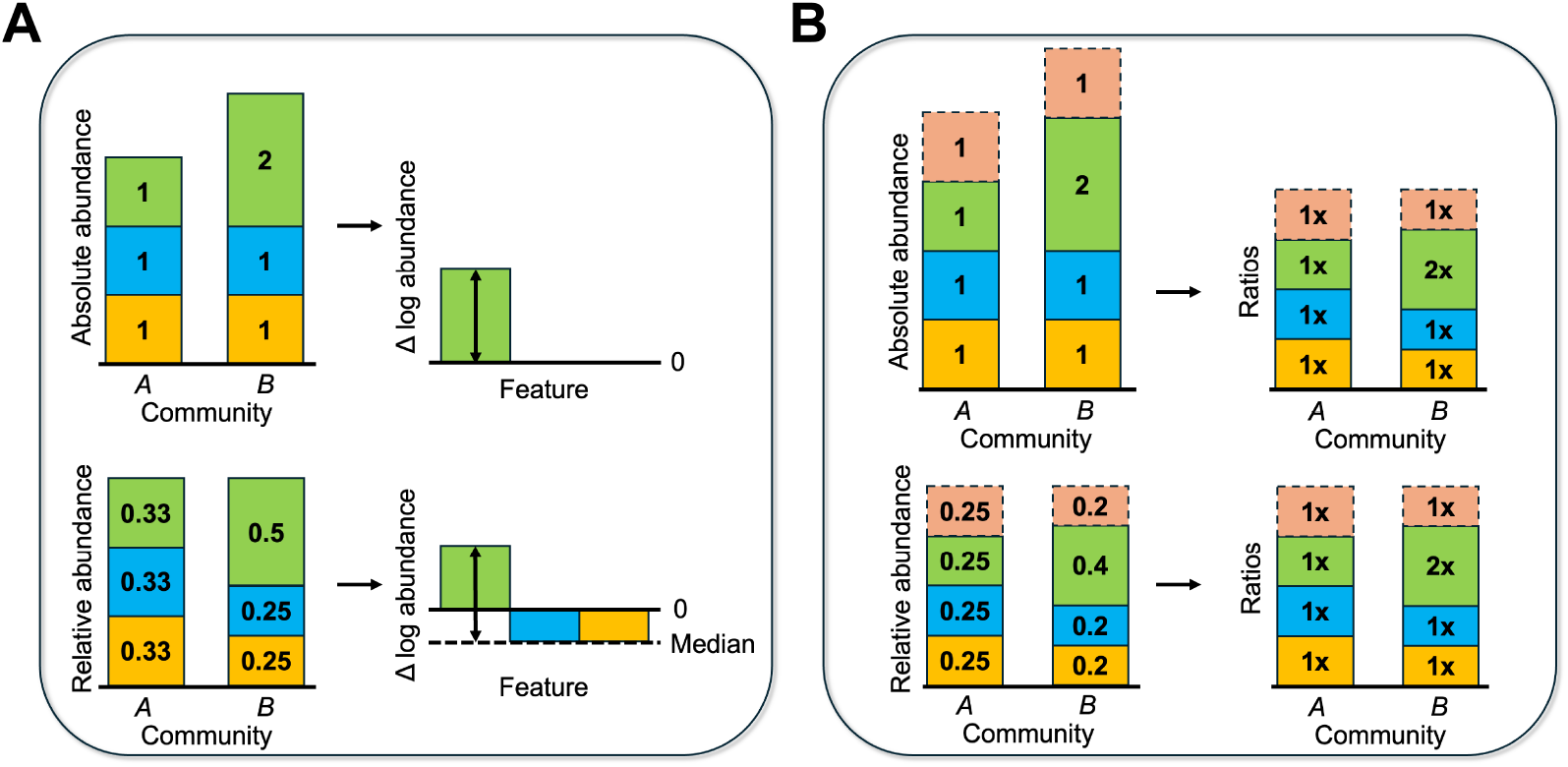
MaAsLin 3 can account for compositionality with or without experimental protocol modifications. **A**. When only relative abundances are available, regressions are first fit on the relative abundance data, and the resulting coefficients are compared to the median of coefficients for each metadatum (median across the features). **B**. When spike-in abundances are known from the experimental protocol, the relative abundances are scaled to the spike-in to compute ratios. These ratios are then used in the regressions. Alternatively, the relative abundances can be scaled by a measure of total community abundance, producing estimated absolute abundances that are then used in the regressions.

**Supplementary Figure 2:**
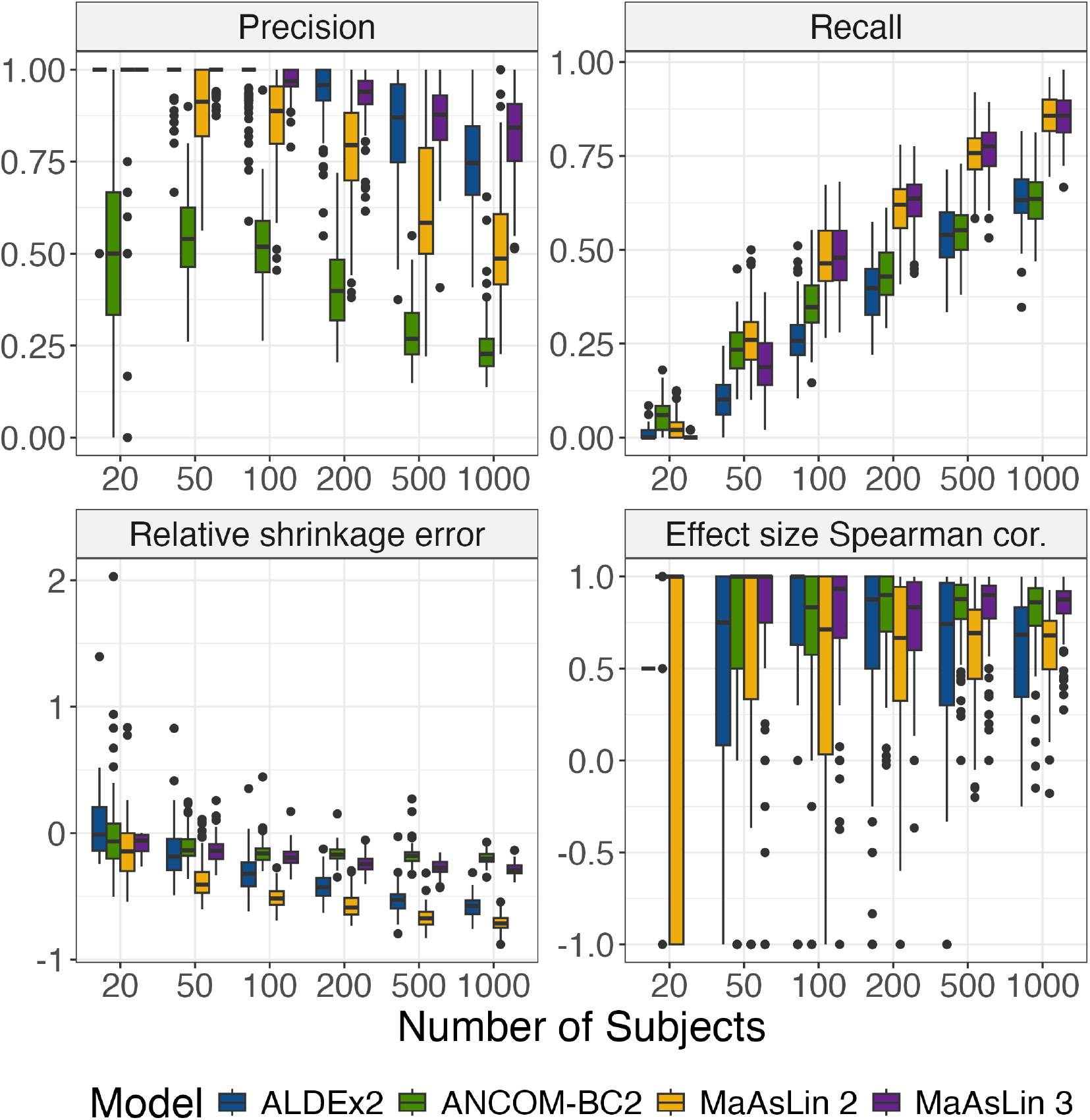
MaAsLin 3 improves accuracy over other DA methods, particularly with high sample sizes. MaAsLin 3 and other common DA methods were run on the 100 synthetic log-normal datasets 2 from **Fig. 1C**. Each metric was calculated as before. 1 is optimal for all metrics except shrinkage, for which 0 is optimal. Each point represents a simulated dataset.

**Supplementary Figure 3:**
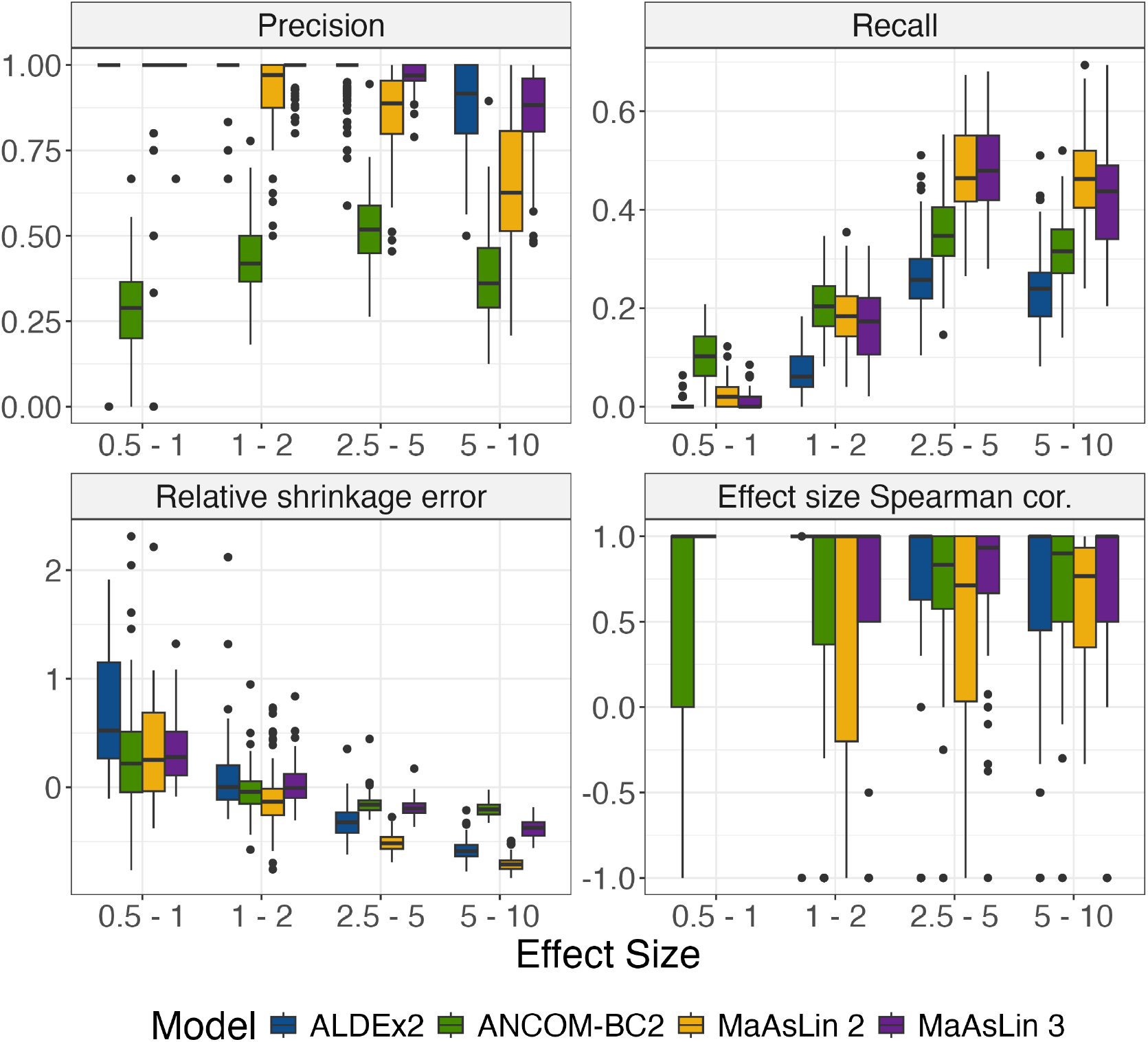
MaAsLin 3 maintains high precision and accurate effect size estimation over a range of biologically relevant effect sizes. MaAsLin 3 and other common DA methods were run on 100 synthetic lognormal datasets from SparseDOSSA 2. The datasets were generated as in **Fig. 1C** but with 100 samples for all datasets and varying effect sizes. Each metric was calculated as before. 1 is optimal for all metrics except shrinkage, for which 0 is optimal. Each point represents a simulated dataset.

**Supplementary Figure 4:**
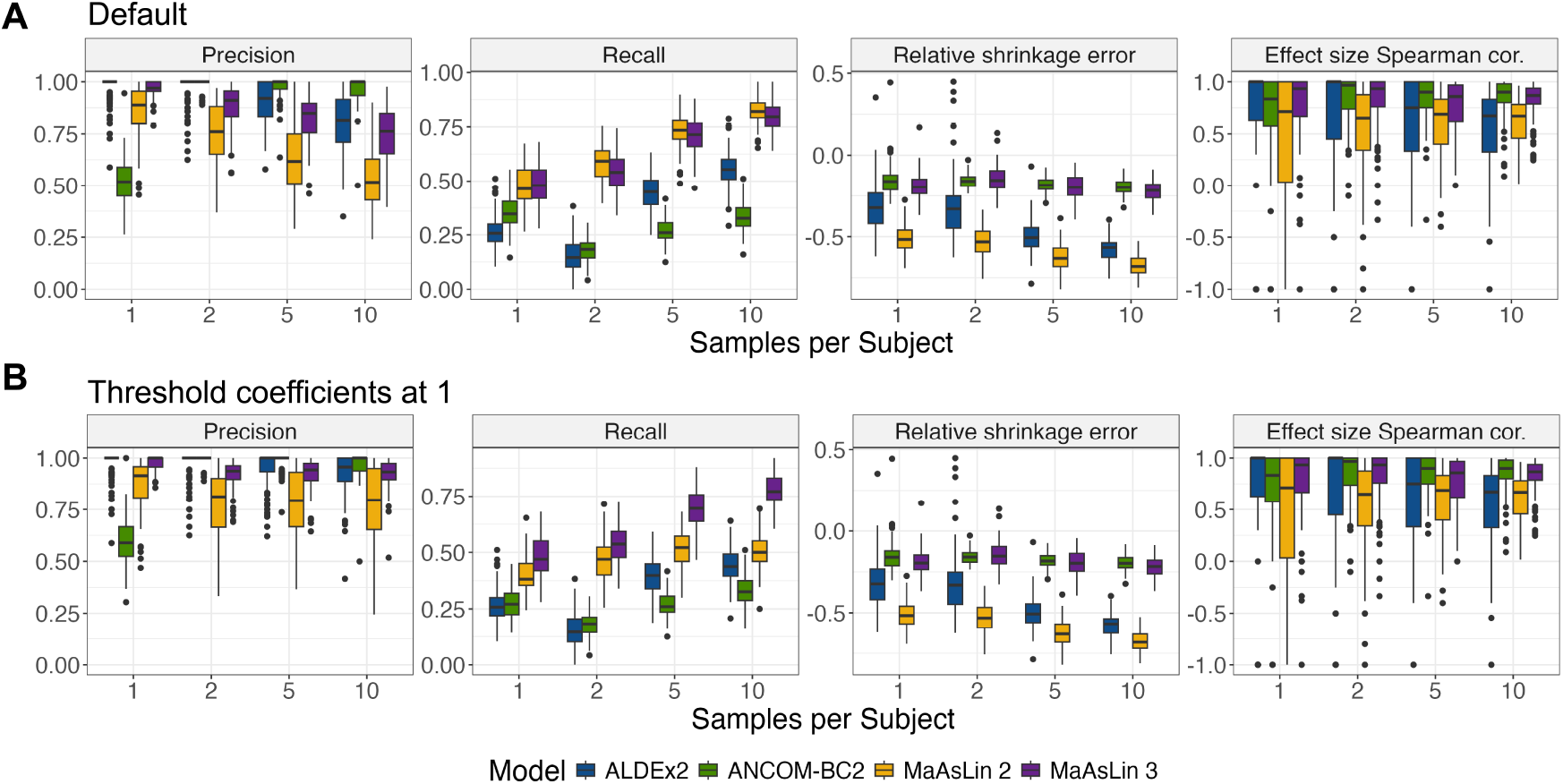
MaAsLin models show high recall with repeated sampling at the cost of reduced precision, though precision can be substantially improved for MaAsLin 3 by thresholding fit coefficients. MaAsLin 3 and other common DA methods were run on 100 synthetic log-normal datasets from SparseDOSSA 2. Each dataset was generated as in **Fig. 1C** but with 100 subjects and varying numbers of samples per subject. For each feature, each subject was given a random intercept drawn from a normal distribution when generating the data. Each point represents a simulated dataset. The metrics were calculated as before on either all associations (**A**) or only associations with fit coefficients larger than 1 in absolute value (**B**). 1 is optimal for all metrics except shrinkage, for which 0 is optimal.

**Supplementary Figure 5:**
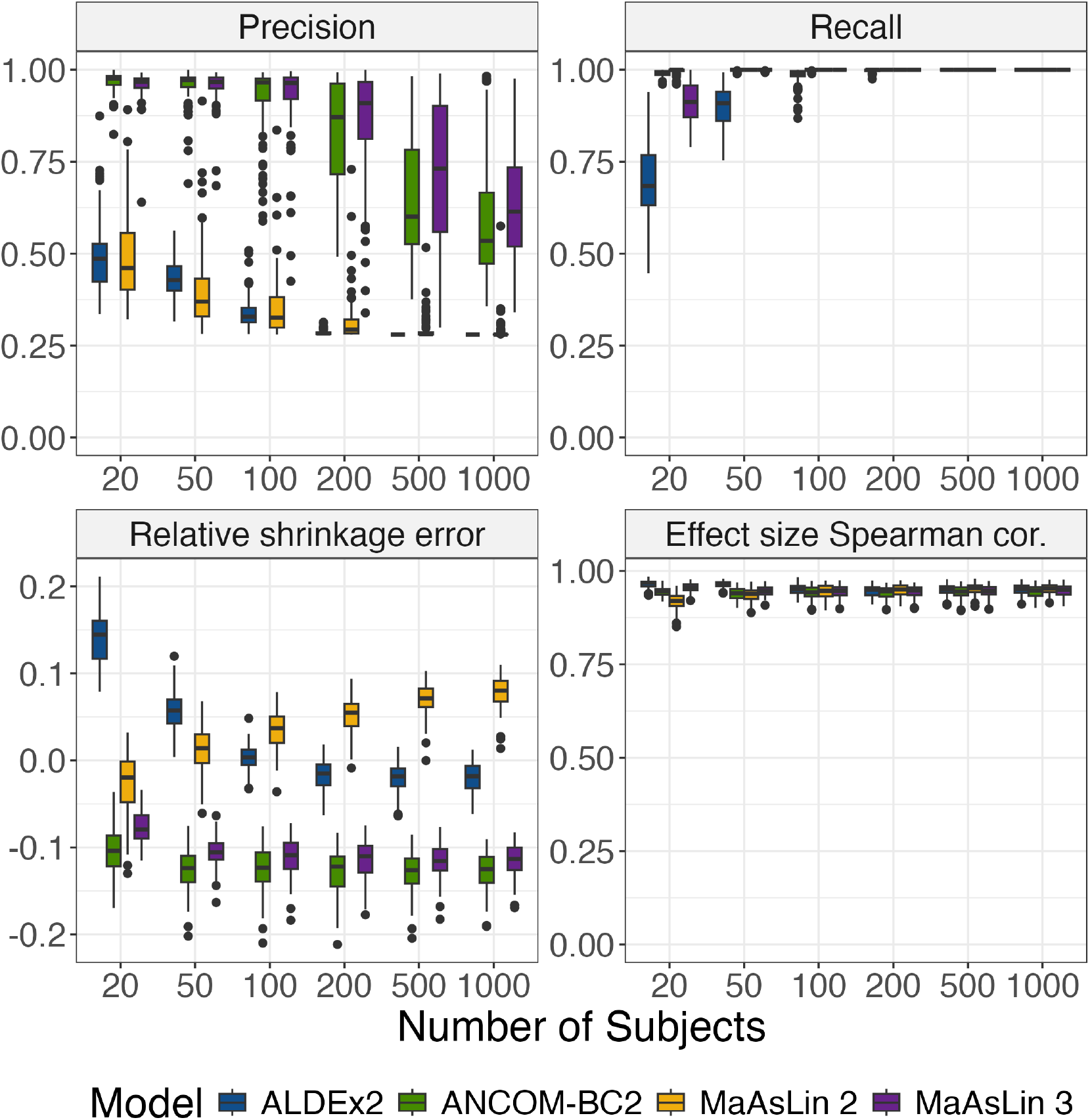
MaAsLin 3 maintains or improves accuracy even when its modeling assumptions are violated. MaAsLin 3 and other common DA methods were run on 100 synthetic datasets generated with the ‘soil’ option of the ANCOM-BC evaluation. For these simulations, 1000 features and 2 groups were simulated with 10% of the feature-metadatum pairs having true associations with coefficients uniform from 2.5 to 5, half of which were positive and half of which were negative. Additionally, 20% of the features were set to have structural zeros in which all samples from one group lacked the feature. Highly skewed and unbalanced read depths (analogous to 16S read count) were drawn using the ANCOM-BC evaluation procedure with a mean depth of 50,000. Metrics were computed as before. 1 is optimal for all metrics except shrinkage, for which 0 is optimal. Each point represents a simulated dataset.

**Supplementary Figure 6:**
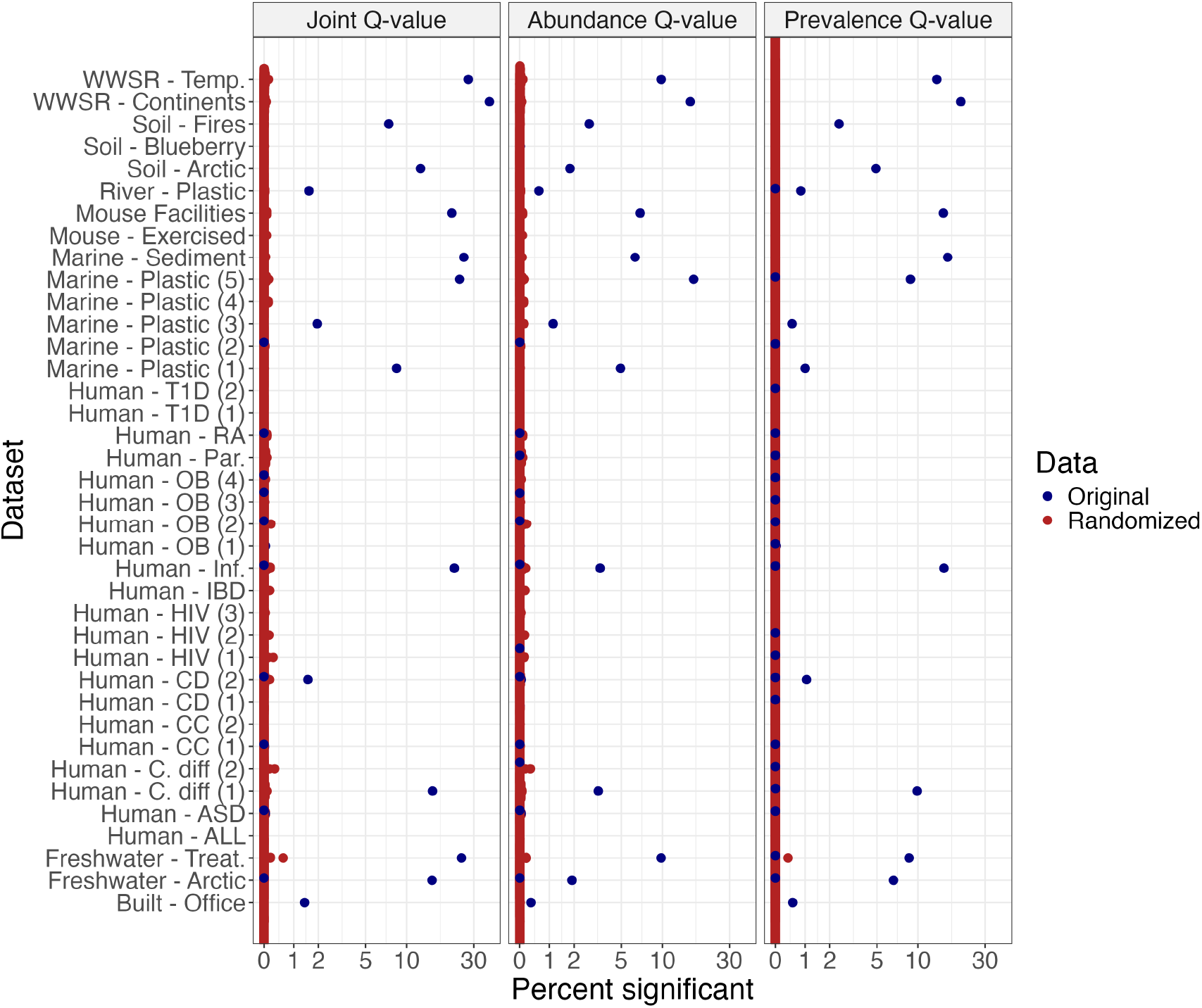
A randomization test using real data shows that MaAsLin 3 almost never produces false positives when all associations are null. Using previous datasets,^1^ 100 mock datasets were created for each real dataset by permuting the binary metadata labels. MaAsLin 3 was then run on the resulting randomized metadata and ASV tables, which should have no associations. For comparison, MaAsLin 3 was also run on the original datasets without randomization, which should have associations if they exist. The percent of significant feature-metadatum associations for both schemes is displayed. 0 is optimal for all randomized settings.

**Supplementary Figure 7:**
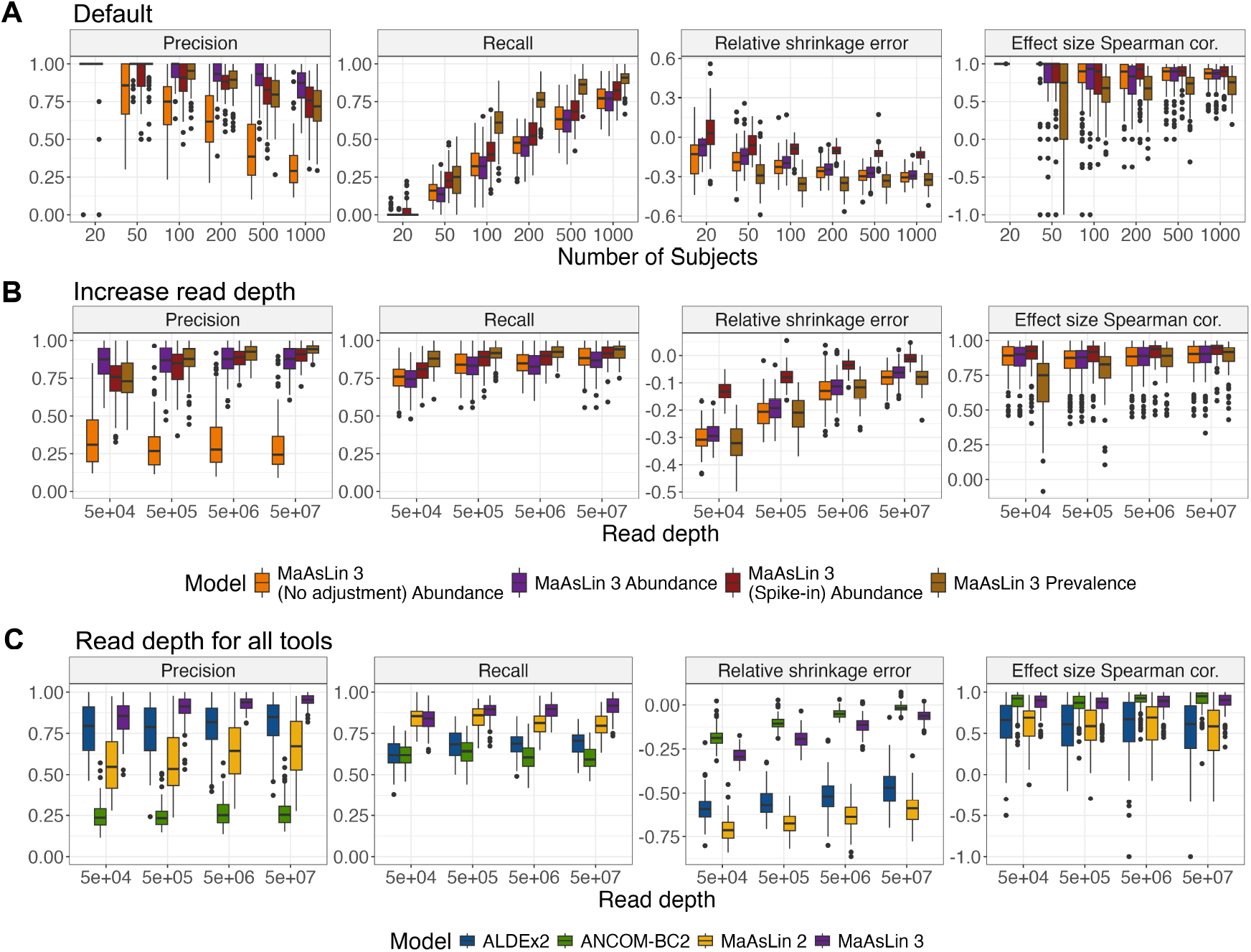
Precision loss with high power can be mitigated by increasing read depth. **A**. Using 100 synthetic log-normal datasets from SparseDOSSA 2, MaAsLin 3 was run with no median adjustment for compositionality (abundance only), with the default adjustment (abundance only), with synthetic spike-in information (abundance only), and with the default prevalence setting. For **A**, the same datasets were used as for **Fig. 1C**. Significant associations (individual q-value less than 0.1) were only considered correct if they matched the true associations in the feature, metadatum, and type of association (prevalence/abundance). The relative shrinkage error and effect size correlation were computed as before. 1 is optimal for all metrics except shrinkage, for which 0 is optimal. Each point represents a simulated dataset. **B**. Datasets were generated as in A but holding the number of subjects fixed at 1000 and varying the mean read depth. **C**. The same metrics were evaluated for all methods on the datasets from B. In this evaluation, significant associations (q-value less than 0.1, joint q-value for MaAsLin 3) were considered correct if they matched the true associations in the feature and metadatum. A mismatch in association type—abundance versus prevalence—was allowed for all methods since no methods besides MaAsLin 3 report association type.

**Supplementary Figure 8:**
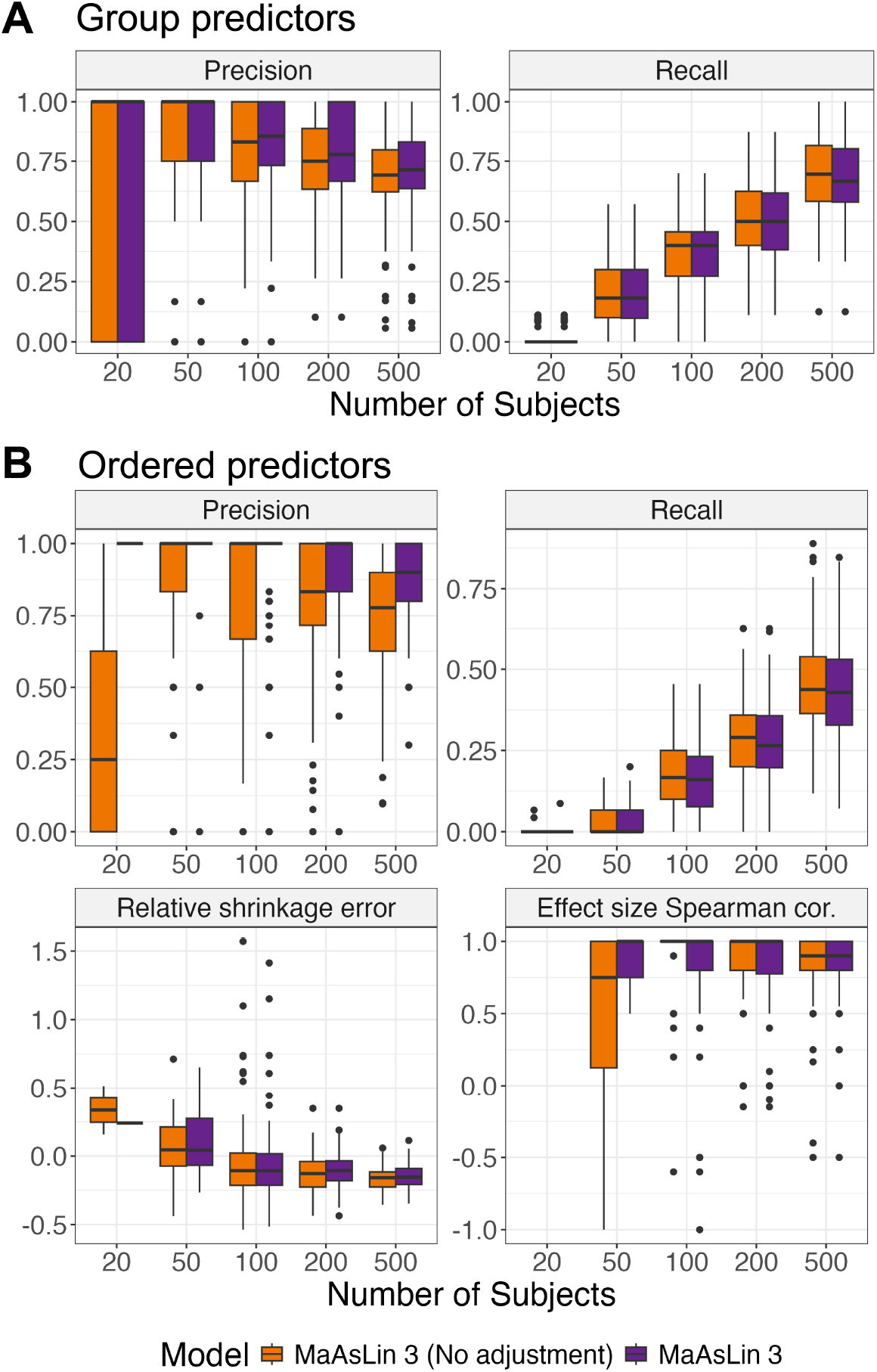
MaAsLin 3 enables inference for group-wise differences and ordered predictors. MaAsLin 3 was run on 100 synthetic log-normal datasets from SparseDOSSA 2. For these simulations, 100 features and 2 metadata (one continuous or binary and one a categorical variable for group-wise or ordered predictors) were simulated with 10% of the feature-metadatum pairs having true associations. Coefficients for the continuous or binary variable were chosen from 2.5 to 5 uniformly. For the group-wise and ordered predictors, a value was chosen from 2.5 to 5 uniformly to represent the most extreme group’s (level’s) difference from baseline, and this value was subdivided according to a Dirichlet(1) distribution to obtain the coefficients for the other groups (levels). Half of the associations were positive; the rest were negative. Half were abundance associations; the rest were prevalence associations. The read depth (analogous to 16S read count) per sample was drawn from a log-normal distribution with a mean of 50,000. Significant associations (q-value less than 0.1, joint q-value for MaAsLin 3) were considered correct if they matched the true associations in the feature, metadatum, and type of association (prevalence/abundance). The metrics were calculated as before. 1 is optimal for all metrics except shrinkage, for which 0 is optimal. Each point represents a simulated dataset. Only the group predictor (**A**) or ordered predictor (**B**) variables were evaluated for accuracy. MaAsLin 3 was run with and without the median compositionality adjustment in both cases.

**Supplementary Figure 9:**
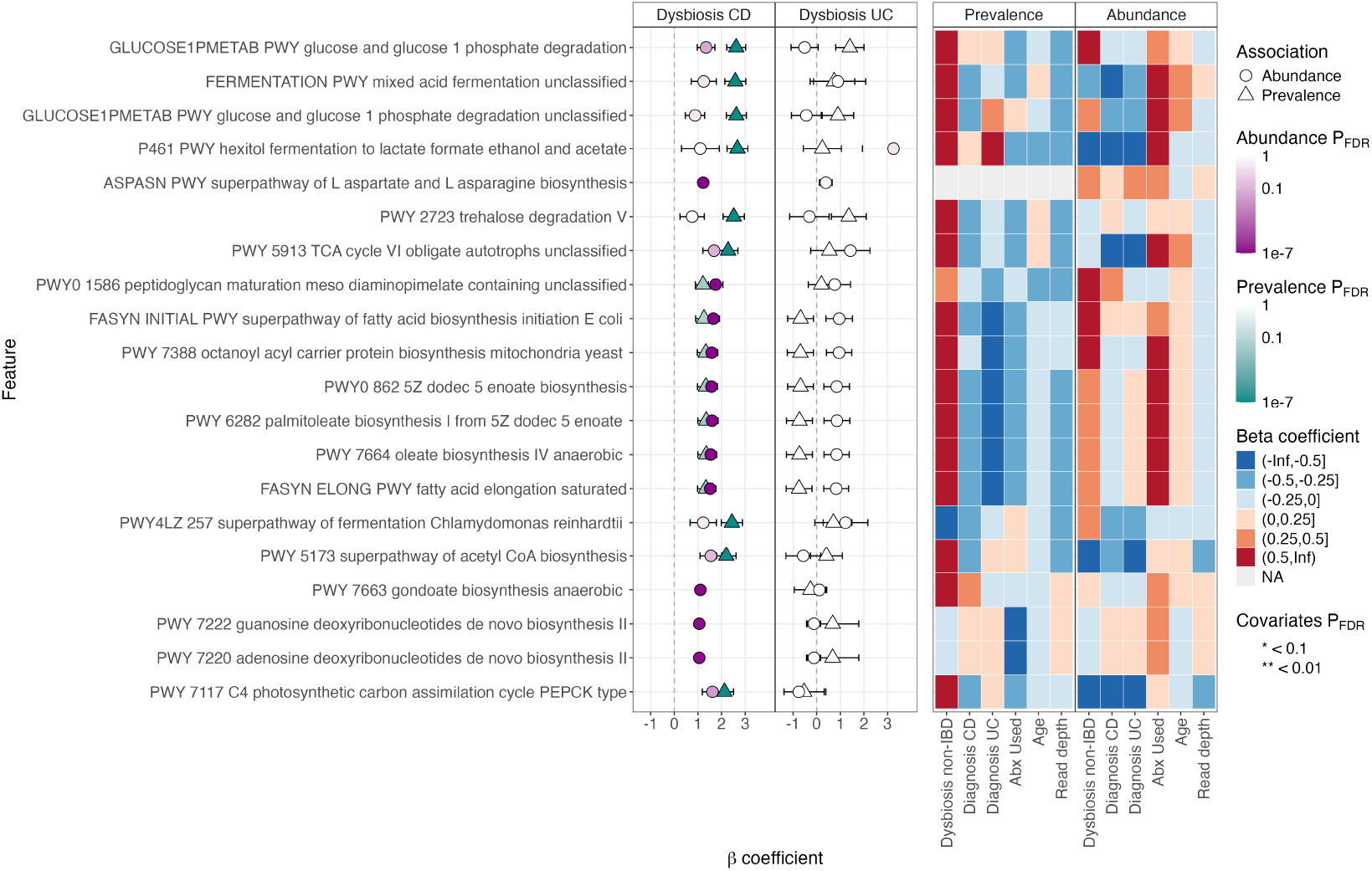
MaAsLin 3 applied to the HMP2 metatranscriptomics data verifies and extends previous findings. The metatranscriptomics pathway abundances from the HMP2 cohort were regressed in MaAsLin 3 using a model that incorporated disease-stratified dysbiosis, disease diagnosis, antibiotic usage, age, read depth, a per-participant random intercept, and the pathway’s metagenomic abundance as a feature-specific covariate.

**Supplementary Figure 10.**
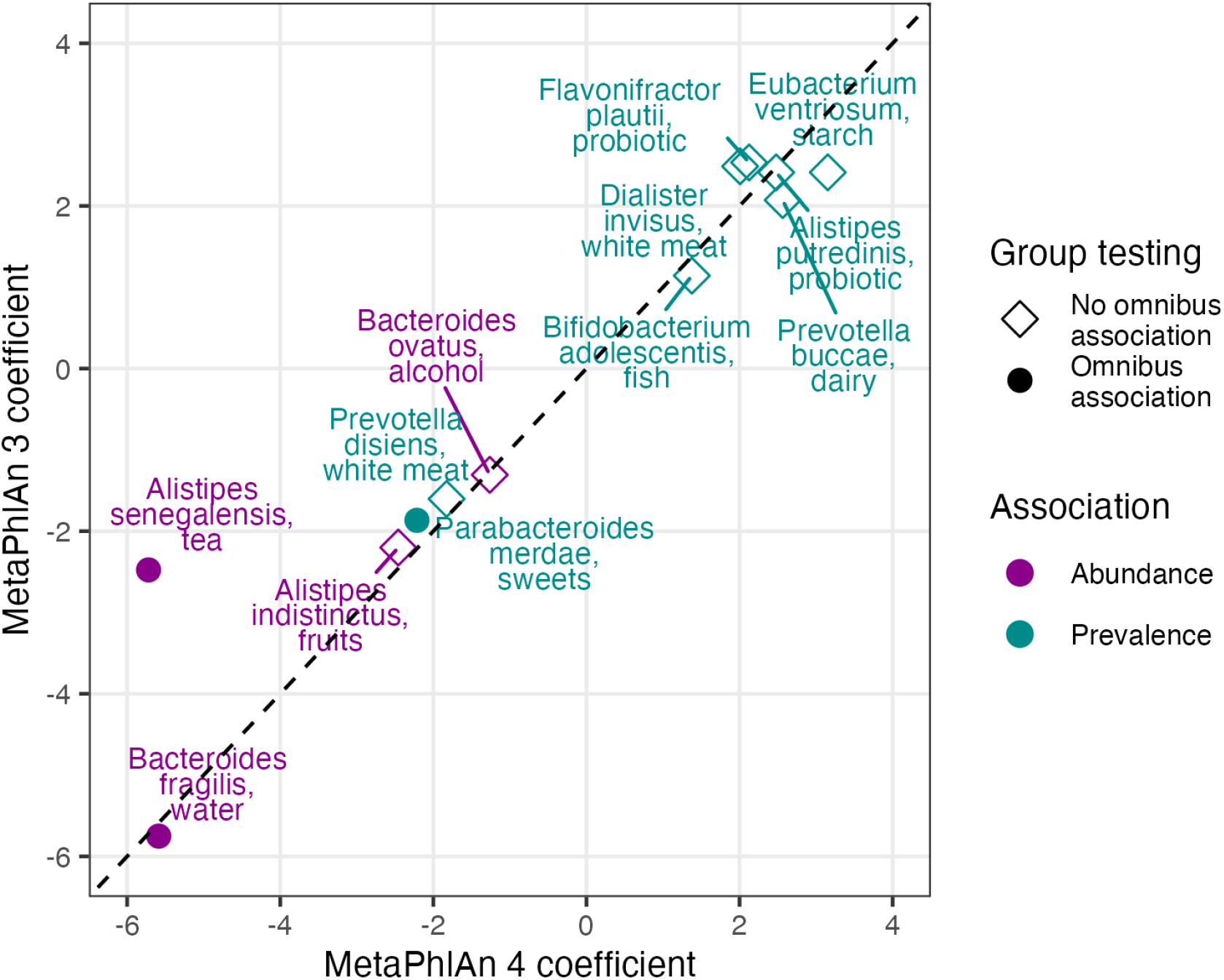
Significant associations in both MetaPhlAn 3 and MetaPhlAn 4 profiles largely agree in fit coefficients. Using the subset of HMP2 participants with CD, abundances were regressed in MaAsLin 3 using a model that incorporated categorical dietary frequency information as a group or ordered predictor along with dysbiosis, antibiotic usage, age, read depth, and a per-participant random intercept. Associations identified as significant using both the MetaPhlAn 3 and MetaPhlAn 4 profiles are plotted.

**Supplementary Figure 11:**
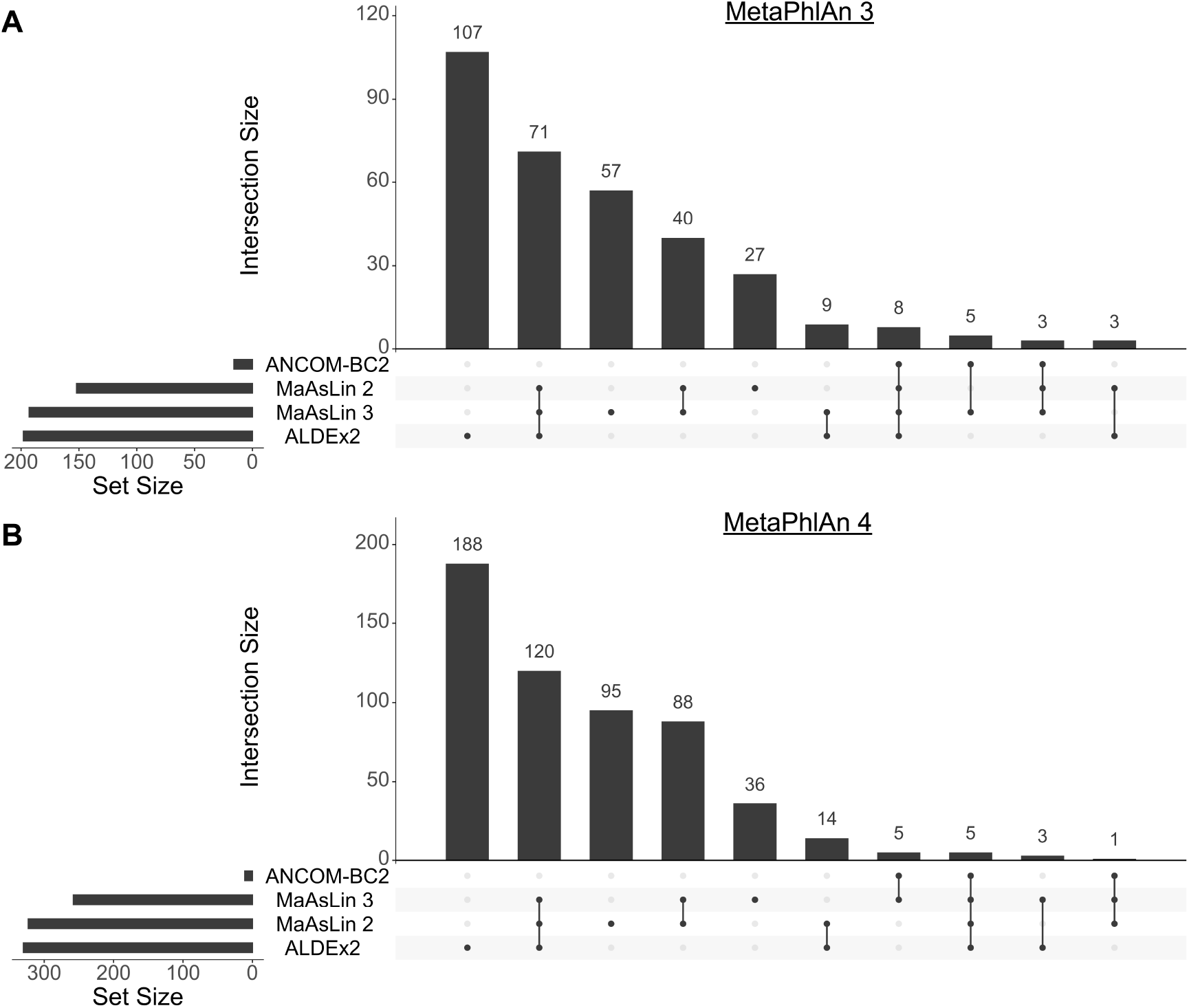
Most IBD associations discovered by MaAsLin 3 overlapped with other methods. The species-level abundances from the HMP2 cohort profiled with MetaPhlAn 3 (**A**) or MetaPhlAn 4 (**B**) were regressed in each method using a model that incorporated disease-stratified dysbiosis, disease diagnosis, antibiotic usage, age, read depth, and a per-participant random intercept (or, for ALDEx2, a fixed intercept subsequently removed from analysis). Because of the possibility for false positives identified in the simulations, only significant (q-value ¡0.1) coefficients with absolute values greater than 1 were evaluated for their overlap.

